# The impact of multiband and in-plane acceleration on white matter microstructure analysis

**DOI:** 10.1101/2023.09.24.559215

**Authors:** Zhengwu Zhang, Arun Venkataraman, Martin Cole, Tianrui Ye, Deqiang Qiu, Feng V. Lin, Benjamin B. Risk

## Abstract

Accelerated imaging has been broadly adopted in diffusion MRI studies, yet little is known about its impacts. Acceleration can achieve higher spatial and q-space resolution in shorter time, reduce motion artifacts, and reduce patient burden. However, it leads to noise amplification, and its impacts in clinical cohorts are poorly understood. This study examined the impact of multiband (also called simultaneous multislice, or SMS) and in-plane acceleration (IPA, also called phase acceleration) in diffusion imaging in forty older adults differing in cognitive status. We evaluated a total of 400 scans from five acquisitions: no acceleration (S1P1); SMS=3 with no in-plane acceleration (S3P1); SMS=3 with IPA=2 (S3P2); S6P1; and S6P2. The number of diffusion directions and b-values was kept constant such that acquisition times varied from 21:28 to 3:56. We found that diffusion metrics were highly sensitive to acceleration factor, with a trend towards higher fractional anisotropy (FA) and lower orientation dispersion (OD) with acceleration. The differences between accelerated and unaccelerated acquisitions could be partly explained by the noise amplification (g-factor) and reduced motion. Intraclass correlations (ICCs) of FA and OD in white matter were excellent in both S1P1 and S3P1 (median >0.8), good but lower in S3P2 and S6P1 (medians around 0.70), and poor to fair in S6P2 (medians 0.46 and 0.57). In-plane acceleration decreased ICC, including areas of high susceptibility distortion. In a comparison of mild cognitive impairment versus healthy controls, acceleration tended to reduce group differences, particularly in the fornix, with greater costs in OD than FA. Our results provide guidance regarding the costs of acceleration (possible biases and reduced effect sizes) while also characterizing the benefits (reduced motion, good reliability at higher multiband with no in-plane).

## 1. Introduction

Diffusion MRI (dMRI) is a non-invasive imaging technique for probing the structure of white matter (WM) in the brain [Le Bihan et al., 1986; Taylor and Bushell, 1985; Merboldt et al., 1985]. WM microstructure features (e.g., fractional anisotropy, neurite orientation and dispersion density imaging metrics [Zhang et al., 2012]) from dMRI play critical roles in understanding neurological and psychiatric disorders [Basser and Pierpaoli, 2011; Le Bihan, 2003; Moseley et al., 1991; Park and Friston, 2013; Pasternak et al., 2018]. However, advanced dMRI can be slow to acquire because it requires the measurement of a large number of volumes along multiple diffusion directions (q-space). To overcome this limitation, MRI acceleration methods have been widely adopted. These methods fall into two main categories: slice acceleration, also called simultaneous multislice (SMS) or multiband [Lee et al., 2005; Feinberg and Setsompop, 2013; Moeller et al., 2010], and in-plane acceleration (IPA), also called parallel imaging, generalized autocalibrating partially parallel acquisitions, phase acceleration, or integrated parallel acceleration techniques [Pruessmann et al., 1999; Griswold et al., 2002]. SMS imaging utilizes a multiband radiofrequency pulse to excite multiple slices at the same time, resulting in a decrease in acquisition time by a factor of *N*, where *N* is the number of slices excited at once. IPA accelerates the acquisition of a single slice by skipping lines of k-space in the phase-encoding direction, with the advantage of reducing echo time and image distortion in addition to a moderate reduction of scan time in a dMRI scan.

Accelerating dMRI acquisition can have a significant impact on image quality and the derived microstructure features. On the one hand, accelerated imaging allows for higher spatial and q-space resolution within a limited time period, which can improve WM microstructure estimates. Shorter acquisitions may also reduce motion. Motion causes artifacts, including bias in derived features [Yendiki et al., 2014; Baum et al., 2018]. Shorter acquisition times can reduce patient burden, an important benefit in clinical scans. IPA reduces geometric distortions due to susceptibility artifacts and results in less blurring [Bammer et al., 2002]. On the other hand, accelerated acquisition can lead to noise amplification, adversely impacting imaging quality. Studies on the costs and benefits of acceleration methods in dMRI are still limited, and various choices of SMS and IPA factors are used in existing brain imaging studies. For example, the Alzheimer’s Disease Neuroimaging Initiative 3 (ADNI3) uses SMS=1 and IPA=2 in their basic acquisition and SMS=3 and IPA=1 in the advanced acquisition [Weiner et al., 2017]. The Human Connectome Project (HCP) young adult study [Uğurbil et al., 2013], UK Biobank [Miller et al., 2016], and ABCD study are using SMS=3 with no IPA [Casey et al., 2018]. The HCP aging and development studies are using SMS=4 [Harms et al., 2018], and NKI Rockland used SMS=4 [Nooner et al., 2012]. Many other smaller studies include phase acceleration (IPA=2) [Tax et al., 2019]. The goal of this study is to document the impact of SMS and IPA on WM microstructure features, with an emphasis on clinical populations.

There are multiple costs affecting image quality in accelerated imaging. First, acceleration induces noise amplification (decreased signal-to-noise ratio) from the statistical errors introduced during the prediction of the single-slice data from the multiband packets [Risk et al., 2018]. The noise amplification can be quantified using the coil geometry factors (*g*-factor) [Breuer et al., 2009]. G-factors vary spatially and tend to be higher in areas where the coil sensitivities are lower and have overlapping profiles, such as subcortical regions and some prefrontal cortical regions [Todd et al., 2017; Risk et al., 2021]. Additionally, IPA results in a 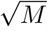 penalty on the signal-to-noise ratio, where *M* is the IPA factor, due to the skipped lines of k-space. The shorter repetition time (TR) in accelerated imaging also results in a reduced signal due to the decreased steady-state longitudinal magnetization.

Resting-state and task-based fMRI studies suggest that moderate SMS factors are helpful in areas of low-to-medium noise amplification, while no acceleration may be optimal in regions with high noise amplification. In Todd et al. [2017], the SNR and t-statistics in a low *g*-factor area (visual cortex) benefited from acceleration but declined in an area of high *g*-factor (ventral medial prefrontal cortex). Moderate acceleration factors (e.g., SMS=4) can improve estimates of task activatation and functional connectivity [Bhandari et al., 2020; Demetriou et al., 2018; Cahart et al., 2022]. However, in many subcortical regions, single-band acquisitions have larger effect sizes than SMS acquisitions and acceleration can substantially degrade activation and functional connectivity studies [Srirangarajan et al., 2021; Risk et al., 2021; Cahart et al., 2022].

Although the costs and benefits of accelerated imaging have been well studied in functional MRI, the impacts in dMRI have received less attention. Zhao et al. [2015] and Duan et al. [2015] conducted a study examining the reproducibility of diffusion tensor imaging (DTI) metrics in eleven healthy participants with SMS=4. They reported fair to good reliability, as measured by the intraclass correlation coefficient (ICC). Furthermore, they compared their ICC values with those of a different study that used single-band data, suggesting better ICC values for the multiband data. Lucignani et al. [2021] evaluated neurite orientation dispersion and density imaging (NODDI) metrics with test-retest data in 8 healthy adults and 9 children with various clinical diagnoses with SMS=2, and a new population with 8 healthy adults and 6 children with SMS=3. The authors concluded SMS=3 was significantly more reliable than SMS=2. Bouyagoub et al. [2021] found some NODDI metrics (orientation dispersion and neurite density) had high reproducibility but disagreement between acquisitions in eight healthy adults across single-band, SMS=2, and SMS=3, where all acquisitions used IPA=2.

Although these previous studies have been important for examining acceleration impacts, key questions remain. First, none of these studies evaluated faster acquisitions, e.g., SMS=6, which may be particularly helpful for reducing patient burden in clinical populations. Second, the studies involved small samples. In some cases, they compared ICCs across different participants, with possible confounding with clinical diagnoses. Third, the effect of increased noise and decreased motion in accelerated imaging on WM microstructure has not been examined. Fourth, the impact on between-group differences has not been examined; for example, diseased versus healthy.

In this study, we analyzed a total of four hundred scans collected during two visits from 20 participants with mild cognitive impairment (MCI) and 20 healthy older participants. In each visit, we collected diffusion data under five acceleration factors: no acceleration (SMS=1, IPA=1) and two SMS factors (3 and 6) crossed by no IPA (IPA=1) and IPA=2. A key aspect of our test-retest design is that the number of diffusion directions is equal across acquisitions, which allows an examination of the costs of noise amplification and the benefits of reduced motion.

Our test-retest and two-sample design allows us to address the following gaps in the literature in quantifying the impact of acceleration in diffusion MRI: 1) extensive comparison of multiple SMS factors (1, 3, 6) and IPA (1, 2) using a test-retest study with large sample size; 2) examining bias and reproducibility of voxel-wise diffusion metric measures (e.g., DTI and NODDI metrics) caused by acceleration; 3) understanding the costs of noise amplification and benefits of motion reduction on bias across accelerations; and 4) comparison of the effect size under different accelerations.

## 2. Data and preprocessing

### 2.1. Participants and imaging method

A healthy older group (n=20, 12 females, ages 62-83) and MCI group (n=20, 12 females, ages 60-86) were recruited from the University of Rochester and its surrounding areas. All participants were consented and the study was approved by the University of Rochester Institutional Review Board. Table 1 summarizes the demographics of the participants.

**Table 1:**
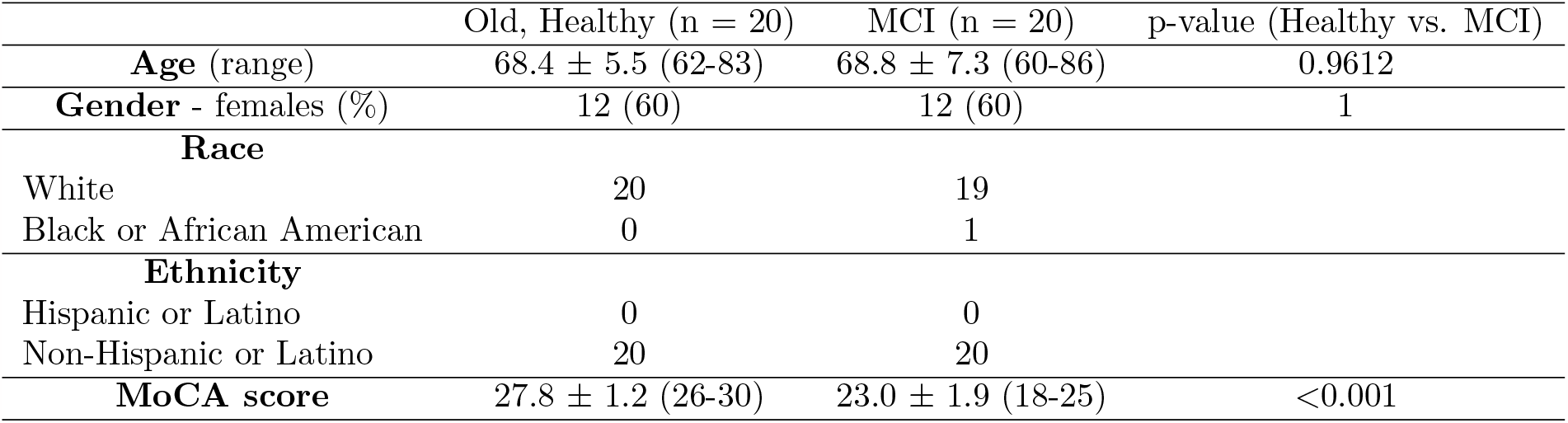
Demographic table for study participants. Age, MoCA scores are represented as mean *±*1 standard deviation with range reported in parentheses. Gender is listed as the number of female participants followed by its percentage in parentheses.

|  | Old, Healthy (n = 20) | MCI (n = 20) | p-value (Healthy vs. MCI) |
| --- | --- | --- | --- |
| **Age** (range) | 68.4 ± 5.5 (62-83) | 68.8 ± 7.3 (60-86) | 0.9612 |
| **Gender** - females (%) | 12 (60) | 12 (60) | 1 |
| **Race** |  |  |  |
| White | 20 | 19 |  |
| Black or African American | 0 | 1 |  |
| **Ethnicity** |  |  |  |
| Hispanic or Latino | 0 | 0 |  |
| Non-Hispanic or Latino | 20 | 20 |  |
| **MoCA score** | 27.8 ± 1.2 (26-30) | 23.0 ± 1.9 (18-25) | <0.001 |

#### Inclusion and Exclusion Criteria

For the healthy older cohort, inclusion criteria were age 60-89, English speaking, adequate vision, self-report free of dementia or MCI, raw total Montreal Cognitive Assessment (MoCA) *≥*26, Rey Auditory Verbal Learning Test (RAVLT) *≥* 50% of the age-adjusted normal value, Geriatric Depression Scale (GDS) *<* 6, and right-handed. For the MCI group, inclusion criteria were the same, except for self-reported memory difficulties and MoCA score between 18 *−*25 and RAVLT *≤*50% of the age-adjusted normal value. In addition, if the participants were on any memory medication (e.g. cholinesterase inhibitor and/or memantine), only patients with constant dosage for at least 3 months before enrollment were included. Exclusion criteria for all groups included neurologic or vascular disorders, documented episode of active psychiatric disorder within the past five years, schizophrenia, living in assisted living or nursing homes, and general MRI contraindications (e.g. claustrophobia, metal implants, pacemaker).

#### Imaging Protocol

The MRI data was collected on a Siemens 3T Prisma scanner (Erlangen, Germany) with a 64-channel head and neck coil. We collected both T1-weighted (T1w) and dMRI data from each participant in two identical sessions separated by up to two weeks. At each session, the imaging sequences included a T1w-MPRAGE (Echo time TE/TR = 2.34*/*2530 ms, TI = 1100 ms, 1*×*1*×*1 mm^3^ resolution, 192 slices/slab) and dMRI consisting of different SMS (multiband acceleration factor) and IPA (iPAT, or integrated phase acceleration techniques, also called GRAPPA factor) settings. Table 2 shows a summary of these protocols. We use notation S*n*P*m* to denote different dMRI protocols, where *n* refers to the SMS factor, and *m* to the IPA factor (e.g., S3P2 denotes SMS factor of 3 and IPA factor of 2). TEs were matched for S3P1 and S6P1 (TE=93 ms) and S3P2 and S6P2 (TE=82.2 ms). All dMRI data were collected using anterior-posterior (AP) phase encoding along with a corresponding reverse posterior-anterior (PA) phase-encoding map for distortion correction. The dMRI scan order was randomized to avoid confounding factor of protocol order (mainly for the consideration of motion). The same 2*×*2*×*2 mm^3^ resolution was used for all dMRI acquisitions with the same 142 diffusion weighting directions (14 interspersed b=0 images, 26 b=1000, and 102 b=2000). The leak-block kernel was used in the reconstruction. Note the in-plane acceleration does not result in large temporal acceleration because the TR is largely dependent on the diffusion preparation time.

**Table 2:** Diffusion MRI acceleration parameters and acquisition time. The 2*×*2*×*2 mm^3^ resolution was used for all acquisitions with 142 diffusion weighting directions (b-values: 14 interspersed b=0, 26 b=1000, and 102 b=2000).

| SMS (S) | IPA (P) | TE/TR (ms) | Acquisition Time (min:sec) |
| --- | --- | --- | --- |
| S1 | P1 | 82/9190 | 21:28 |
| S3 | P1 | 93/3378 | 8:10 |
| S3 | P2 | 82.2/2740 | 7:05 |
| S6 | P1 | 93/1723 | 4:17 |
| S6 | P2 | 82.2/1404 | 3:56 |

### 2.2. Image processing

#### Diffusion MRI Preprocessing

We used FSL topup to correct for susceptibility distortions with the reverse phase-encoding map [Andersson et al., 2003]. Following distortion correction, FSL eddy was used to correct for motion and eddy current effects [Andersson and Sotiropoulos, 2016] with outlier images replaced using the Gaussian process (GP) algorithm in Andersson et al. [2016].

#### Quality Control

In quality control (QC), we quantified the *absolute motion, signal to noise ratio (SNR)* for b=0 data, and *contrast to noise ratio (CNR)* for b=1000 and b=2000 data. Implementation of QC in [Bastiani et al., 2019] was used to quantify *absolute motion*, SNR for b=0 data, and CNR for b=1000 and b=2000 data. The absolute motion was computed in the following way: each dMRI volume was compared with the first volume and the average displacement was calculated. For each dMRI scan, we obtained one number to quantify the absolute motion in this scan. The SNR and CNR were defined at the voxel level. At each voxel, SNR was computed as the mean of the b=0 data across all acquisitions normalized to the standard deviation over the same acquisitions, i.e.,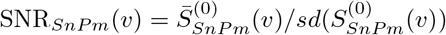where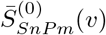 is the mean of all b=0 images at the voxel *v*, and *sd*(*·*) computes the standard deviation. Note we have 14 b=0 images for each acceleration setting, which were all used to compute the SNR. The CNR was used to quantify diffusion-weighted volumes. For a given diffusion direction, the measured diffusion signal is denoted as 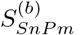, and we used the GP model in the eddy tool [Andersson et al., 2016] to predict the diffusion signal, denoted as 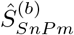. Based on the two quantities, the CNR was calculated as

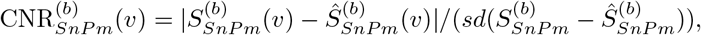

where *sd* 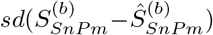 computes the standard deviation of the prediction residuals. We also calculated a normalized CNR as *CNR* 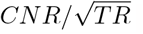 to assess CNR adjusted for TR.

#### T1 Preprocessing

The T1 image was first denoised using a non-local means algorithm [Descoteaux et al., 2008]. Advanced Normalization Tools (ANTs) [Avants et al., 2009] was then used to remove the bias field from the image [Tustison et al., 2010]. Finally, the T1 image was co-registered to the dMRI using ANTs.

#### Generation of WM Mask

A WM mask was generated using the atropos function in ANTs [Avants et al., 2009] applied to the MNI152 T1 1 mm standard image available in FSL. Additionally, the MNI152 T1 image was run through the volbrain pipeline (https://volbrain.upv.es/) to generate a mask of the subcortical gray matter regions. Because the ANTs white matter map also included subcortical gray matter regions, the subcortical gray matter mask was subtracted from the atropos output to generate the final MNI WM mask used for masking and visualization.

#### Diffusion Metrics

To understand the acceleration’s effect on dMRI data analysis, we extracted the following dMRI-derived microstructure metrics.

*Voxel-level diffusion tensor metrics:* After the preprocessing steps, the dipy [Garyfallidis et al., 2014] package was used to fit diffusion tensors to extract tensor metrics, including fractional anisotropy (FA) and mean diffusivity (MD). Only b=1000 data were used for fitting diffusion tensors. In this study, the voxel-level analyses focus on FA since it is one of the most popular DTI metrics.

*Voxel-level NODDI metrics:* The AMICO package (https://github.com/daducci/AMICO) was used to calculate NODDI metrics using the entire multishell data (b=1000 and b=2000). Three NODDI metrics were computed: orientation dispersion (OD) index, isotropic volume fraction (ISOVF), and intracellular volume fraction (ICVF).

The voxel-level features were transformed to MNI space for downstream statistical analyses. The co-registered T1 in the dMRI space was used for each participant to find the warping function to align the diffusion space data to the MNI space.

#### Voxel-wise Motion Quantification

We also quantified the voxel-level motion for each dMRI scan. We enabled the “slice-to-vol” motion correction option in FSL eddy and recorded the motion (translation and rotation) estimated for each batch of MB slices [Andersson et al., 2017]. The translation enables the same motion effect for all voxels, but the rotation generates heterogeneous motion impact on different voxels, i.e., voxels far away from the center will be impacted more by the rotation. Globally, we used absolute motion as the measure of aggregate motion for a scan. Locally, in each voxel, we used the chord length calculated from the mean rotation matrix as the voxel-specific measure of motion.

## 3. Methods

### 3.1. Exploratory data analysis

#### 3.1.1. Motion, SNR, and CNR

0, CNR in b=1000, CNR in b=2000, and adjusted CNRs for each acquisition. For each of these measures, we calculated the Cohen’s d effect size by pooling data from all participants. Specifically, we calculated the mean and standard deviation for each acquisition and then calculated the Cohen’s d between two acquisitions by computing the difference in means divided by the pooled standard deviation.

#### 3.1.2. Noise amplification analysis

We characterized noise amplification using the *apparent g-factor* 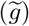 for the acceleration method S*n*P*m* [Preibisch et al., 2015] defined as

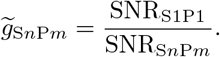

The 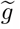-factor was computed for each (S*n*P*m*, S1P1) pair in each session and then averaged in the MNI space.

### 3.2. Possible biases in FA and OD

#### 3.2.1. Mean FA and OD maps for each acquisition

We examined the impact of acceleration on voxel-wise FA and OD measures. We calculated the mean FA and mean OD for different accelerations using all scans in the MNI space. The ICBM DTI-81 atlas [Oishi et al., 2008] with 48 WM regions (shown in Figure S.4) was used to summarize FA and OD measures in different WM regions for a region-based comparison. We used all scans to create box plots from the FA and OD values for each region in the atlas.

#### 3.2.2. Examining the role of g-factor and motion on possible biases

The differences between acquisitions are “possible biases” since we don’t have a gold standard. By definition, S1P1 has no noise amplification and hence does not suffer from noise-amplification-induced biases. On the other hand, S1P1 also has increased absolute motion due to longer scan times, which can introduce bias. Accelerated acquisitions may have increased bias from noise amplification but decreased bias from reduced motion. We examined the mean difference between the accelerated (S*n*P*m*) and unaccelerated images, i.e., S*n*P*m* -S1P1.

To examine the influence of increased noise and reduced motion on FA and OD, we predicted S*n*P*m* -S1P1 for FA or OD at voxel *v* in MNI space for *v* = 1, …, *V* . The predictors were the mean 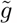-factor (calculated across images) and mean motion difference caused by rotation at voxel *v* (S1P1-S*n*P*m*), denoted as *x*_*v*1_ and *x*_*v*2_, respectively. The motion caused by translation was not considered since it produces the same motion at all voxels, and therefore its impact is contained in the global mean of S1P1 -S*n*P*m*. Taking the S3P1 data as one example, we utilized all S3P1 data from 40 subjects and randomly assigned each subject to one of the three categories, training, validation, and testing, at a ratio of 0.6:0.2:0.2, respectively. This splitting ensures the independence between training, validation, and testing datasets. For the training data, we computed three mean maps in MNI space: mean S*n*P*m*-S1P1 map, mean 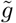-factor map, and mean rotation difference map. Similar maps were computed for both validation and testing sets. The training maps were used to train our predictive models using the following methods: linear regression, generalized additive model(GAM), random forest, XGBoost, support vector machines, and gradient boosting. We also considered transformations and interactions: 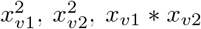 log(*x*_*v*1_), and log(*x*_*v*2_). Throughout the training, the models’ hyperparameters and predictors (including the transformations) were selected according to their performance on the validation set. Including 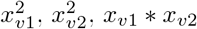, *x*_*v*1_ *\* x*_*v*2_ and log(*x*_*v*1_) did not improve the predictive performance and therefore their results were not reported. Only log(*x*_*v*2_) was found to be helpful with the prediction, and its results were reported. We also studied using only *x*_*v*1_ or *x*_*v*2_ as the predictor to assess the relative roles of motion versus acceleration-induced noise amplification. Once each model was finalized, we evaluated its performance on the testing data using the correlation between predicted and actual data.

### 3.3. Reproducibility analysis

The reproducibility relates to the robustness of a measurement/feature and its discriminative ability. A feature will have high reproducibility if it can distinguish different participants with less variability in repeated measures. Leveraging the repeated measures of different accelerated images, we studied the reproducibility of diffusion metrics using intra-class correlation coefficients (ICC). We define the ICC using the random effects formulation: 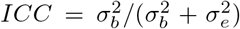where 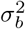 is the between-subject variance and 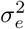 is the within subject variance. This is estimated using (*MSB − MSW*)*/{MSB* + (*k −* 1)*MSW*)*}*, where *MSB* is the mean squared between-subject, *MSW* is the mean-squared within subject, and *k* = 2 in our case [Shrout and Fleiss, 1979]. For FA and OD, we also analyzed the subset of MCI patients, details of which appear in the Web Supplement.

### 3.4. In-plane acceleration and distortion analysis

In-plane acceleration shortens the EPI readout, reducing the geometric distortions while enabling a larger matrix size to be acquired at a given echo time. However, IPA does not result in appreciable temporal acceleration since the TR is largely dependent on the duration taken up by the diffusion preparation in dMRI. For example, in our experiment, S3P1 took 8 mins and 10 seconds, while S3P2 took 7 mins and 5 seconds. We examined whether IPA is helpful for analyzing brain areas susceptible to large distortion. A mask was built based on the distortion maps calculated from the motion outputs of the FSL eddy tool. More specifically, we registered the distortion maps to MNI space and then averaged the registered maps. A threshold of 16 was applied to the absolute value of this average map to extract high-distortion regions. We then studied the reproducibility of FA measured within this mask.

### 3.5. Impact of acceleration on group comparisons

Another important task in brain imaging analysis is to identify differences between groups. We compared how diffusion metrics differed between healthy controls and MCIs under different accelerations. Let *Y*_*irv*_ be the feature of interest (e.g., FA or OD value) for the *i*-th participant (*i ∈{*1, …, 40 *}*) and *r*-th observation (*r ∈{*1, 2 *}*) in the *v*-th voxel. We restricted the analysis to voxels in the WM mask. We computed the adjusted effect size by fitting a linear model with age, gender, session, absolute motion, and group (MCI versus healthy controls) for each voxel. Then the adjusted Cohen’s d was equal to group coefficient (negative indicates lower in MCI) divided by the residual standard deviation. This approach adjusts for possible confounding due to higher motion in the MCI group. We then inspected how Cohen’s d changed across acquisitions. We also computed Cohen’s d using the conventional formula (ignoring motion and other predictors), and the results were similar but generally indicated a larger proportion of voxels with decreased effect sizes in accelerated acquisitions (not reported here).

Next, we used linear mixed models to examine group differences in the fornix, which is a significant WM bundle located in the medial diencephalon and forms part of the limbic system. It plays a crucial role in memory and has garnered attention in recent research into Alzheimer’s disease (AD) [Brown et al., 2017]. A fornix template from Brown et al. [2017] was applied to select the locations for our analysis. We fit a linear mixed model with age, gender, absolute motion, and group (MCI versus healthy controls) including a random intercept for participants. We then studied the p-values associated with the group difference across different acquisitions.

## 4. Results

### 4.1. Exploratory data analysis

We show one coronal slice from a randomly selected participant in Figure 1 after the pre-processing steps in Section 2.2. The top row shows images with intensity normalized within each image (intensity ranges from 0 to 1 in each image) and the bottom row shows the same image but normalized across images (one intensity bar ranging from 0 to 1 for all images). Visually, we can see how acceleration impacts the contrast within each image (top row) and how there is a tendency for the noise to increase as the acceleration increases, which is particularly evident when comparing S6P2 to S1P1 (bottom row). We use arrows to draw the reader’s attention to the brainstem/pons region to show the noise amplification.

**Figure 1.**
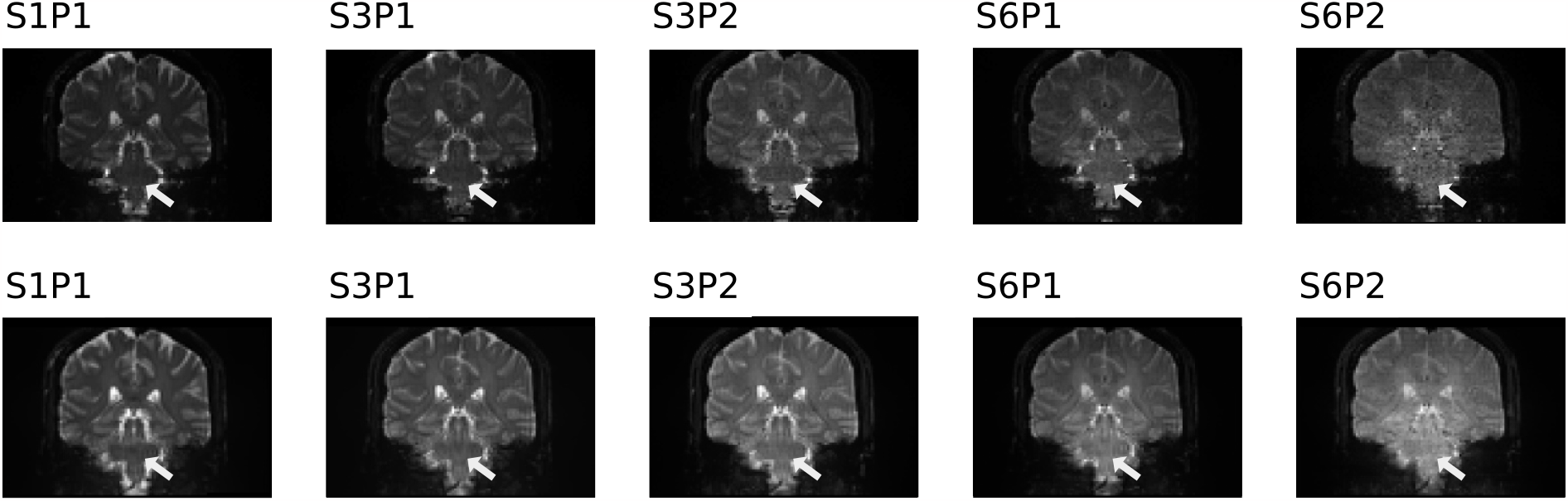
Representative coronal slices from one participant over various acceleration factors after pre-processing. Top row shows data intensity normalized within images (intensity ranges from 0 to 1 in each image). Bottom row shows data with normalized intensity across images (one intensity bar ranging from 0 to 1 for all images). Arrows pointing to brainstem/pons region to show noise amplification related to acceleration.

#### 4.1.1. Motion, SNR, and CNR

QC metrics obtained from the FSL eddy tool are displayed in Figure 2. The numbers on the top of each panel show effect sizes between S1P1 and the accelerated images. In Table S.1, we compared the QC metrics between P1 and P2 accelerations.

**Figure 2.**
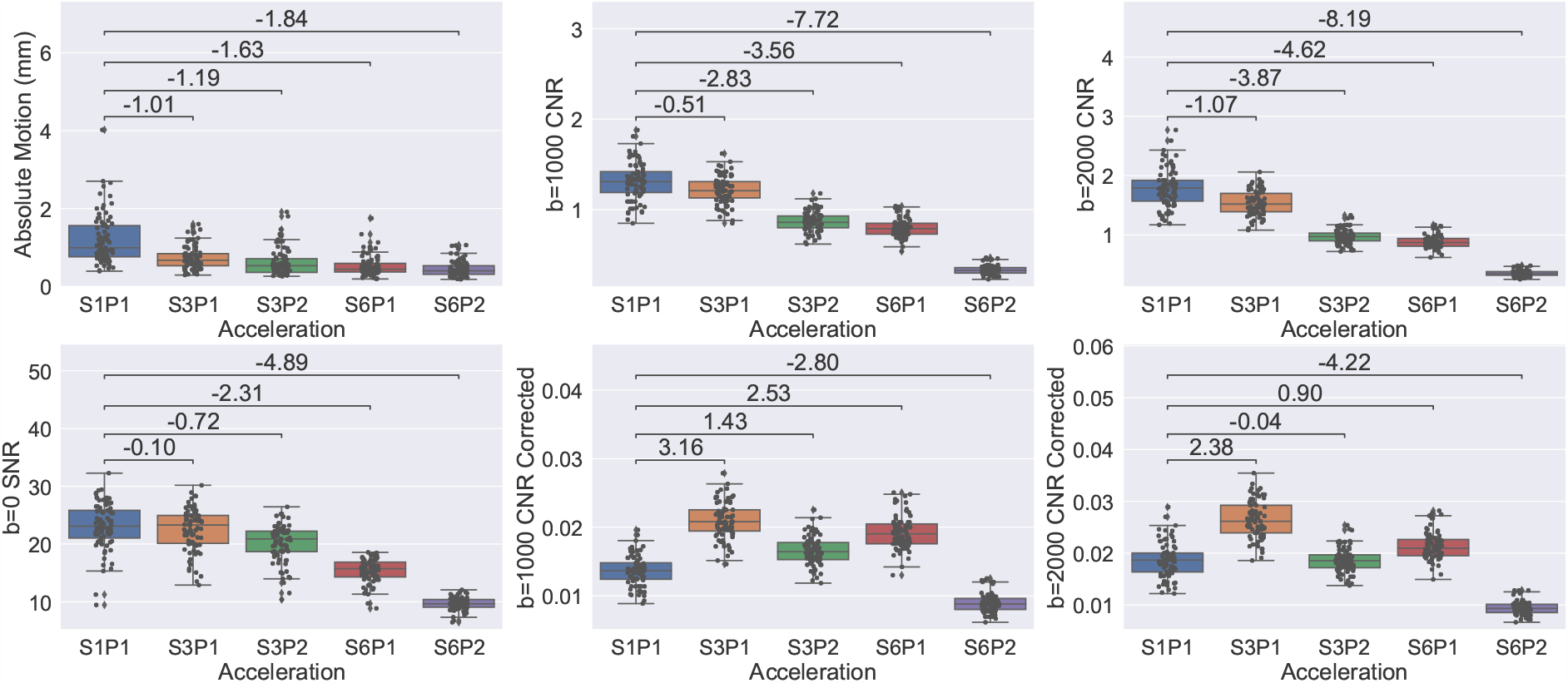
QC metrics of absolute motion (left in row one), SNR (left in row two), CNR and corrected CNR of b=1000 shell (middle column), and CNR and corrected CNR of b=2000 shell (right column). The corrected CNR is equal to 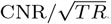. The numbers above bars represent Cohen’s d effect size.

#### Absolute Motion

Absolute motion decreases in accelerated acquisitions. Due to the long acquisition time of S1P1, the effect sizes between the motion in S1P1 and the accelerated acquisitions are large (all Cohen’s d*>*1, Figure 2). The effect sizes comparing the motion in S3P2 to S3P1 and S6P2 to S6P1 are moderate (see Table S.1). In Figure S.1, we separated the healthy controls from the MCIs. On average, MCIs have more head motion than healthy controls, indicating that motion is a confounder when comparing the two groups.

#### SNR and CNR

The SNR decreases with acceleration, but the decrease from S1P1 to S3P1 is relatively small (Cohen’s d = -0.10 with similar medians) compared with the drops between S1P1 and the other accelerated protocols (i.e., S3P2, S6P1 and S6P2, Figure 2). The same trend can be observed for the CNR of b=1000 and b=2000 shells, but with more notable differences between S1P1 and S3P1. CNR is more sensitive to acceleration at higher b-values, as the Cohen’s d for b=2000 are larger than for b=1000. We also adjusted the CNR with respect to the repetition time. Through these adjusted CNRs, we see that per unit time, S3P1 gives the best CNR. Also, from Table S.1 we see that the CNR is highly sensitive to IPA, with very large Cohen’s d for S3P2 versus S3P1 (*< −* 2.5) and S6P2 versus S6P1 (*< −* 7), with greater sensitivity at higher b-values, i.e., the Cohen’s d of the decrease in CNR at the b=2000 shell is larger than at the b=1000 shell.

#### 4.1.2. Noise amplification analysis

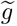**-factor**: Figure 3 shows the averaged 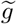-factors computed from b=0 images for each acquisition protocol. We see a general increase in 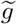-factor with greater acceleration. Frontal areas including regions near the subcallosal cortex have higher 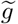-factor compared to posterior areas. Comparing P2 with P1 acquisitions, we see large increases in 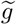-factor in most brain regions (particularly the deep WM regions, the brainstem, and cerebellum). The occipital lobe is less impacted by noise amplification. In fact, some regions in the occipital lobe in S3P1 and S3P2 have 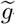-factor values less than *<* 1, indicating higher SNR than S1P1.

**Figure 3.**
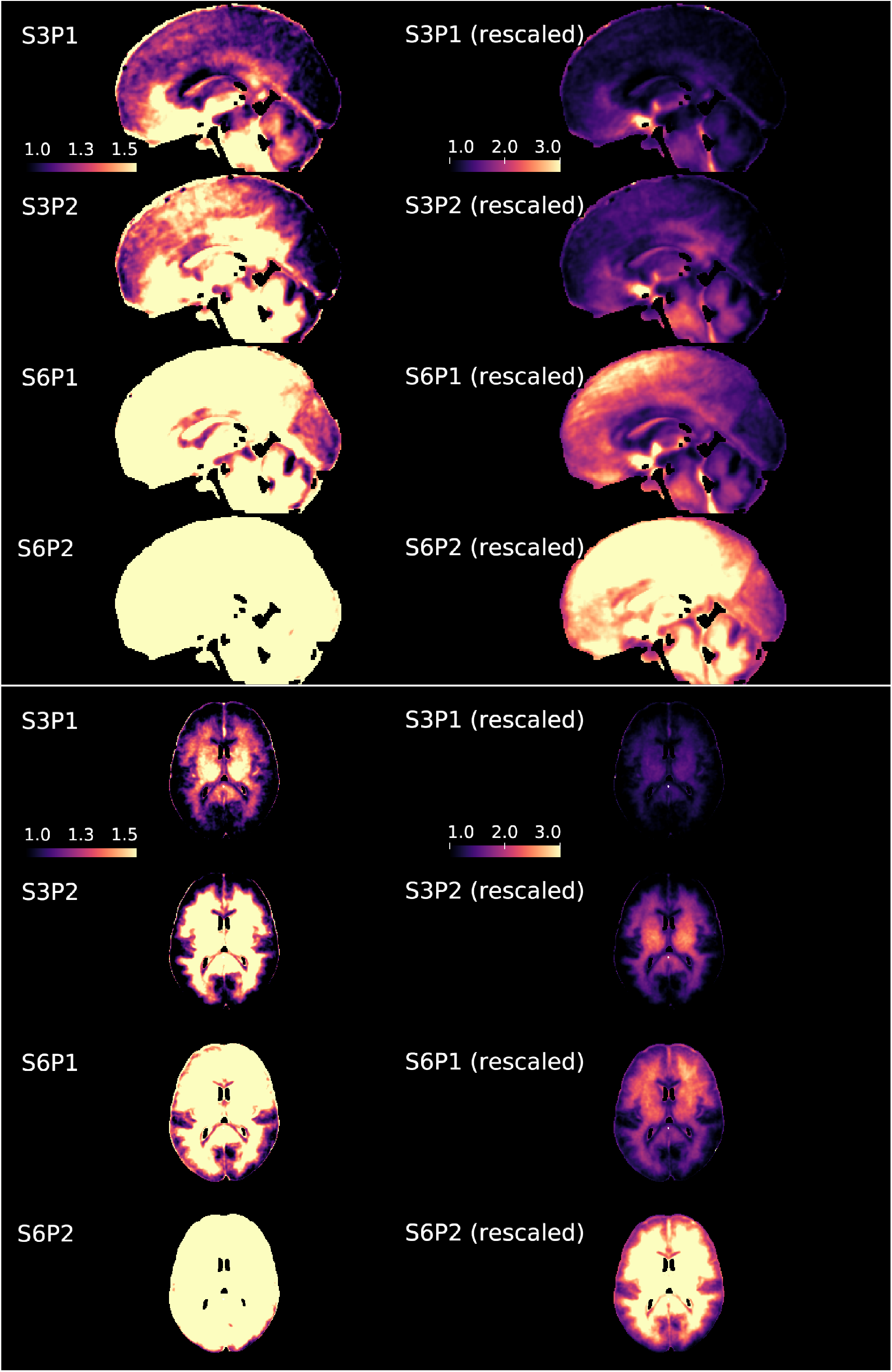
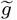-factor maps averaged over all scans. Depicted are sagittal slices (MNI=0) and axial slices (MNI=13). Brighter colors indicate greater noise amplification (poorer image quality).

### 4.2. Possible biases in FA and OD

#### 4.2.1. Mean FA and OD maps for each acceleration

White matter tracts in FA are clearly delineated in S1P1, S3P1, S3P2, and S6P1, but with a trend of increasing FA across the brain with increasing acceleration (top of Figure 4). The cortical gray matter regions exhibit a marked increase in FA in S6P2, resulting in a washed-out appearance. There is a line of higher FA values delineating the boundary between the putamen and pallidum in S1P1 that becomes somewhat more prominent in S3P1 and prominent in S6P1. In S6P2, the divisions between subcortical gray matter regions and white matter tracts are blurred, as many areas have values greater than 0.4. These white matter divisions in S6P2 are visible at an alternative color scale (0 to 0.8), although with less contrast between regions than the other acquisitions (Figure S.2). Box plots of FA values in the first 20 ROI regions defined in the ICBM DTI-81 atlas show clearer differences across accelerations (top of Figure 5) and spatially heterogeneous di across the brain. Box plots of all 48 regions in the atlas can be found in Figure S.3 and Figure S.4 displays the 48 ROIs. White matter tracts in FA are clearly delineated in S1P1, S3P1, S3P2, and S6P1, but with a trend of increasing FA across the brain with increasing acceleration (top of Figure 4). The cortical gray matter regions exhibit a marked increase in FA in S6P2, resulting in a washed-out appearance. There is a line of higher FA values delineating the boundary between the putamen and pallidum in S1P1 that becomes somewhat more prominent in S3P1 and prominent in S6P1. In S6P2, the divisions between subcortical gray matter regions and white matter tracts are blurred, as many areas have values greater than 0.4. These white matter divisions in S6P2 are visible at an alternative color scale (0 to 0.8), although with less contrast between regions than the other acquisitions (Figure S.2). Box plots of FA values in the first 20 ROI regions defined in the ICBM DTI-81 atlas show clearer differences across accelerations (top of Figure 5) with some regions appearing more similar (e.g., region 1, cerebellum white matter) while others differ greatly (e.g., region 6, fornix). Box plots of all 48 regions in the atlas can be found in Figure S.3 and Figure S.4 marks the locations of all 48 ROIs.

**Figure 4.**
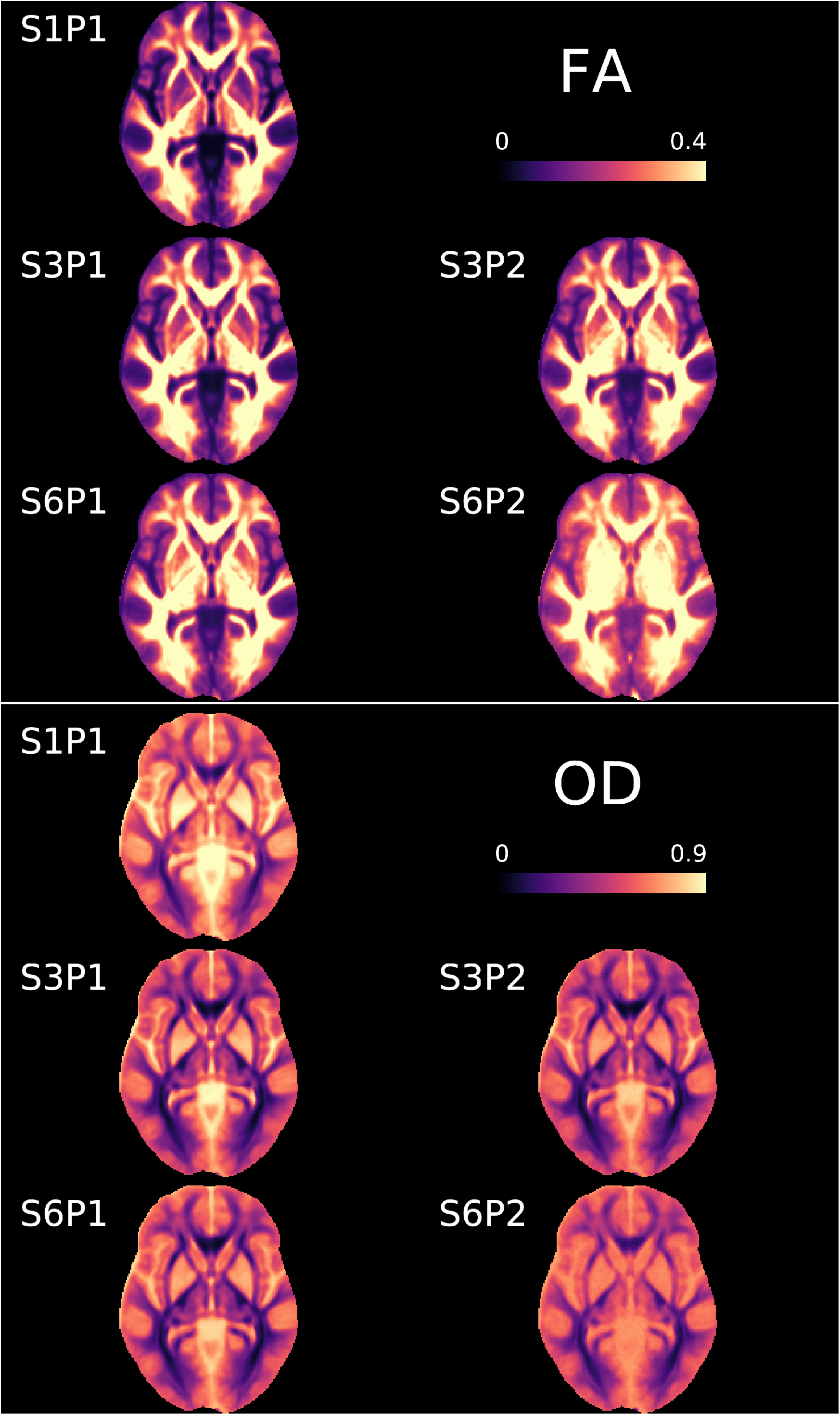
FA (top) and OD (bottom) averaged across all scans.

**Figure 5.**
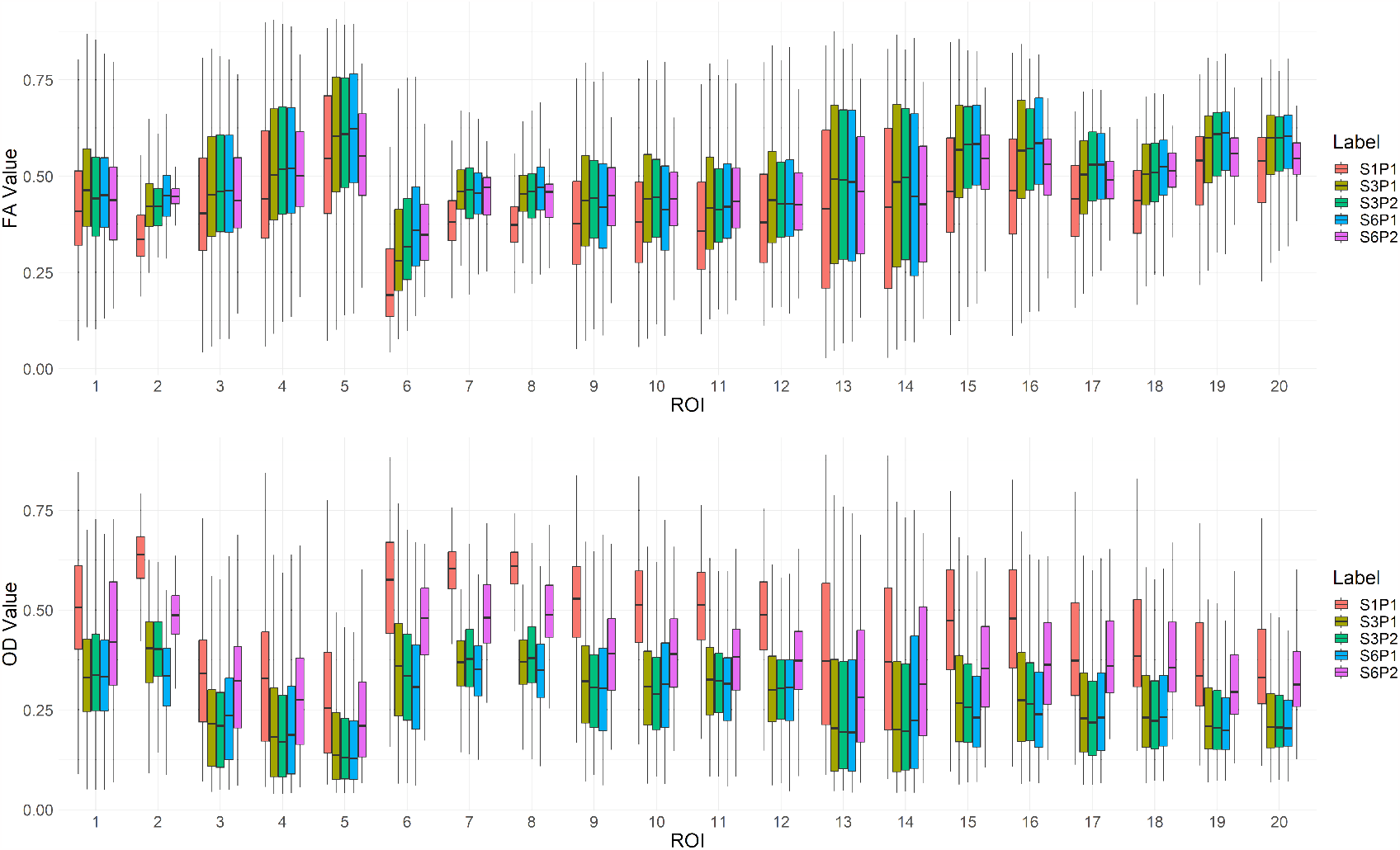
Box plots of FA (top) and OD (bottom) averaged across all scans in the first 20 ROIs in the ICBM-DTI-81 white-matter labels atlas.

OD values trend highest in S1P1 and tend to decrease with acceleration (Figure 4). Overall, patterns trend in the inverse direction of FA, as WM tracts have lower OD values (bottom of Figure 5). S6P2 again has a notably washed-out appearance. Compared with FA, OD seems more significantly impacted by acceleration: the OD in S1P1 is much higher than in accelerated images. For example, region 1 (cerebellum) has higher OD in S1P1 than others, and again we see large differences in region 6 (fornix).

#### 4.2.2. Examining the role of 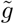-factor and motion on possible biases

Tables 3 presents the S*n*P*m*-S1P1 prediction results using the 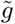-factor (*x*_1_) and motion (*x*_2_). In the main manuscript, we mainly summarize results with no phase acceleration (Table 3), since later we found phase acceleration significantly reduces overall reproducibility (in Section 4.3) and does not help with data analysis in brain regions with large susceptibility distortions (in Section 4.4). For S3P2 and S6P2, see Table S.2. The best prediction performance on the test dataset is *r* = 0.54 for S3P1-S1P1 FA, *r* = 0.41 for S6P1-S1P1 FA, *r* = 0.60 for S3P1-S1P1 OD, and *r* = 0.57 for S6P1-S1P1 OD. We can draw several conclusions from this table:

**Table 3:** Correlation between predicted S*n*P*m*-S1P1 and measured S*n*P*m*-S1P1 in an independent test dataset using 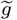-factor (*x*_1_) and motion difference (*x*_2_) for data with no phase acceleration. Phase acceleration results are in Table S.2. The best performance under each scenario is in bold.

| Model | S3P1-S1P1 FA |  |  |  | S6P1-S1P1 FA |  |  |  |
| --- | --- | --- | --- | --- | --- | --- | --- | --- |
| | $(x_1, x_2)$ | $(x_1, x_2, \log(x_2))$ | $x_1$ | $x_2$ | $(x_1, x_2)$ | $(x_1, x_2, \log(x_2))$ | $x_1$ | $x_2$ |
| Linear Regression | 0.51 | **0.54** | **0.52** | 0.30 | 0.29 | 0.38 | 0.27 | 0.25 |
| GAM | **0.53** | 0.54 | 0.51 | 0.32 | 0.36 | 0.37 | 0.29 | 0.26 |
| Random Forest | 0.42 | 0.42 | 0.42 | 0.23 | 0.36 | 0.36 | 0.17 | 0.17 |
| XGBoost | 0.43 | 0.44 | 0.51 | 0.31 | 0.40 | 0.40 | 0.27 | 0.26 |
| SVM | 0.44 | 0.44 | 0.51 | **0.33** | 0.40 | 0.40 | **0.29** | **0.28** |
| Gradient Boosting | 0.51 | 0.50 | 0.50 | 0.32 | **0.41** | **0.41** | 0.28 | 0.26 |

| Model | S3P1-S1P1 OD |  |  |  | S6P1-S1P1 OD |  |  |  |
| --- | --- | --- | --- | --- | --- | --- | --- | --- |
| | $(x_1, x_2)$ | $(x_1, x_2, \log(x_2))$ | $x_1$ | $x_2$ | $(x_1, x_2)$ | $(x_1, x_2, \log(x_2))$ | $x_1$ | $x_2$ |
| Linear Regression | 0.58 | 0.59 | **0.60** | 0.29 | 0.51 | 0.51 | 0.22 | 0.36 |
| GAM | 0.59 | 0.59 | 0.60 | 0.30 | 0.55 | **0.57** | 0.30 | 0.37 |
| Random Forest | 0.55 | 0.55 | 0.50 | 0.22 | 0.52 | 0.52 | 0.20 | 0.32 |
| XGBoost | 0.57 | 0.57 | 0.59 | **0.30** | 0.55 | 0.54 | 0.30 | 0.37 |
| SVM | 0.58 | 0.57 | 0.59 | 0.30 | **0.57** | 0.56 | **0.32** | **0.39** |
| Gradient Boosting | **0.59** | **0.59** | 0.59 | 0.29 | 0.55 | 0.55 | 0.30 | 0.37 |

- The strong prediction performance suggests that the 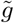-factor and motion can account for a significant portion of the variation in the difference between accelerated and unaccelerated images for both FA and OD.
- In S3P1, we discovered that the 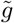-factor more strongly predicts the differences from S1P1 (with *r* = 0.54 in FA and *r* = 0.60 in OD) than motion (with *r* = 0.33 in FA and *r* = 0.30 in OD).
- Comparing S6P1 to S1P1, the prediction performance of 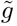-factor and motion are now more similar for FA (*r* = 0.29 and *r* = 0.28, respectively), while motion has stronger predictive performance (*r* = 0.39) than 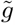-factor (*r* = 0.32) for OD.

### 4.3. Reproducibility Analysis

#### 4.3.1. Voxel-wise diffusion metrics

Figure 6 shows the ICC maps for FA (first column) and OD (third column), and the differences from S1P1 (e.g., S3P1 -S1P1 for the first image in columns 2 and 4). Note here the WM mask from Section was applied. Figure 7 shows box plots of the ICC values in Figure 6. From these voxel-wise ICC values, we can see that

- Both the FA and OD metrics have high reproducibility in the S1P1 and S3P1 protocols (the median ICC values are over 0.8) (Figure 7). Further acceleration substantially reduces the ICC values.
- ICCs of both FA and OD are spatially heterogeneous under all accelerations, with higher values in the occipital and parietal lobes and the corpus callosum and lower values around the brain stem and cerebellum.
- From the ICC difference maps, we see a roughly balanced mix of blue and red on the S3P1-S1P1 map. The blue color dominates after more acceleration (e.g., for S3P2, S6P1, and S6P2).
- Comparing S3P2 with S3P1 and S6P2 with S6P1, we see an overall large decrease in ICC values from the phase acceleration (e.g., in FA, median ICC for S3P1 = 0.84 versus S3P2=0.72; median ICC for S6P1=0.71 versus S6P2=0.46, Figure 7).

**Figure 6.**
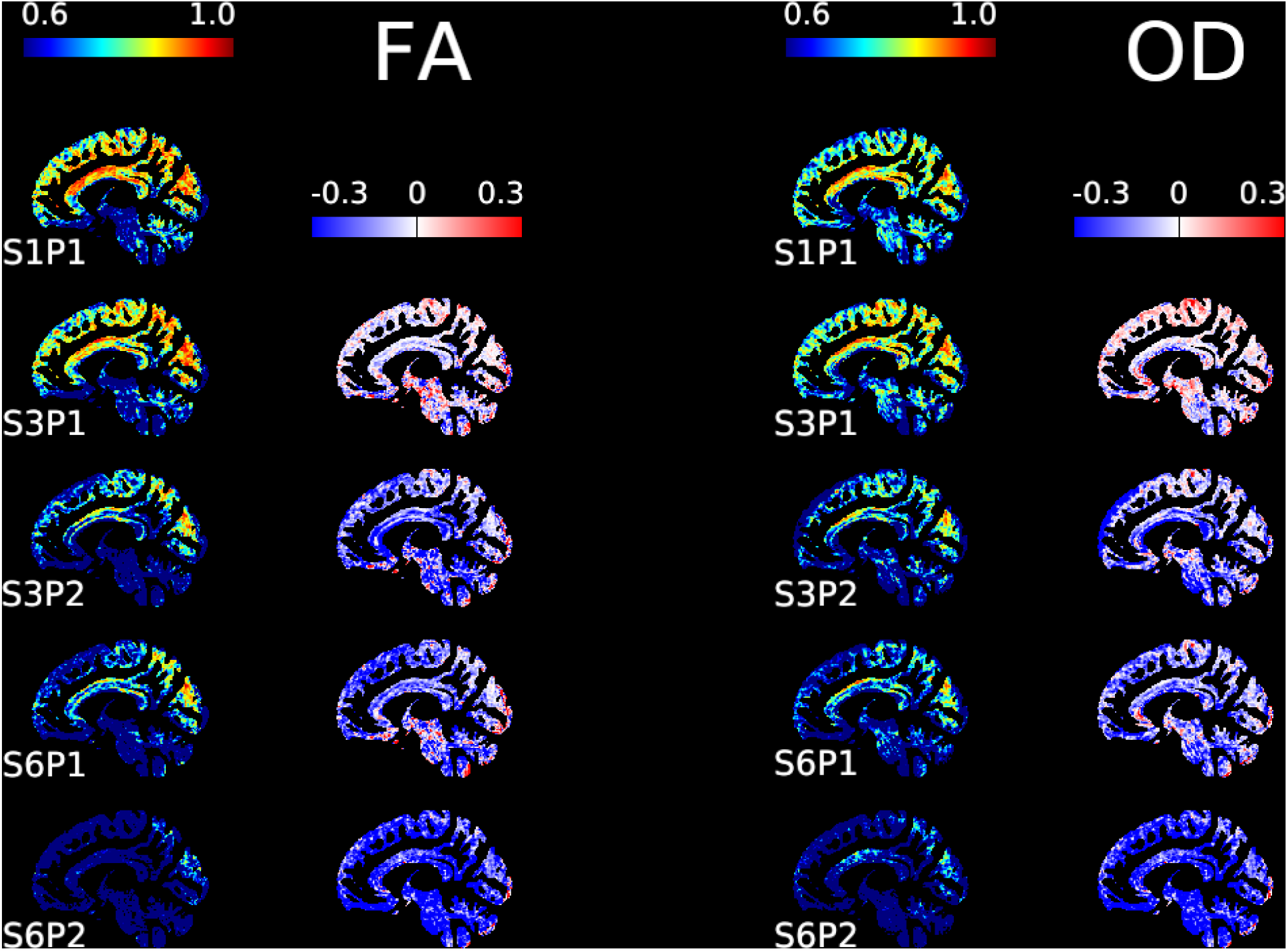
ICC values of FA (left two columns) and OD (right two columns) in white matter with corresponding difference maps (calculated as the difference from S1P1, i.e., S3P1 -S1P1 for the first image in columns 2 and 4).

**Figure 7.**
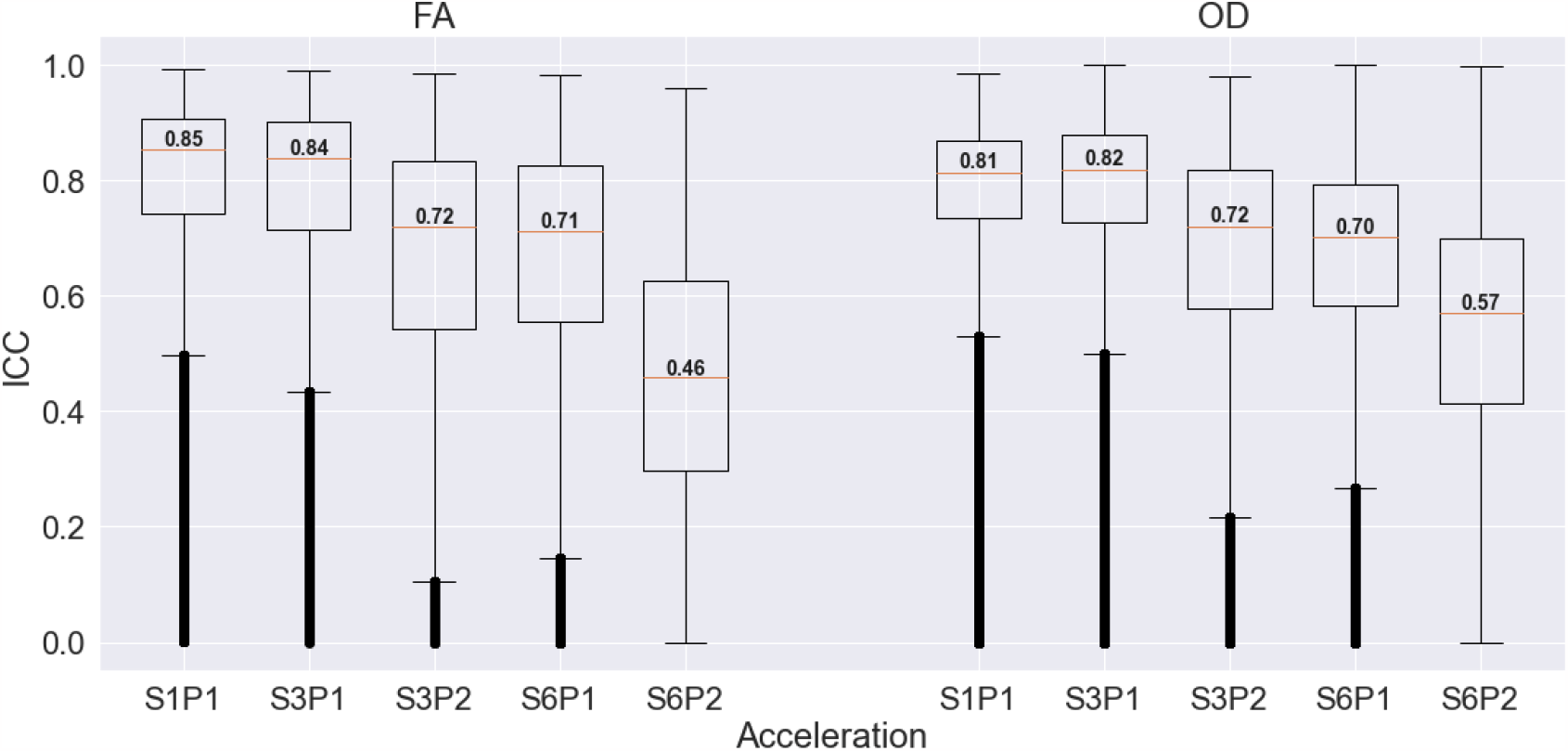
Boxplot of ICC values from voxels shown in Figure 6 for FA (left) and OD (right). Median values for each boxplot are shown in the figure.

We also examined whether the spatial patterns in ICC were similar in the subset of participants with MCI (Figure S.5). The ICC maps for the subset of MCI participants were similar to the ICC maps from all participants.

In the Web Supplement, we describe the ICC findings for ICVF and ISOFV (Figure S.6). We again see S3P1 and S1P1 are similar, with some decreases in ICC with higher acceleration. However, in contrast to FA and OD, in some areas of the brain stem the ICCs are higher than S1P1 for all accelerated acquisitions.

### 4.4. In-plane acceleration and distortion analysis

We analyzed the ICC values in high-distortion regions. In Figure 8 left panel, we compared ICC values for the FA in S3P1 to S3P2 and S6P1 to S6P2. The third row of Figure 8 shows the ICC difference maps, where we see that the red color is dominating, indicating that P1 acquisitions are more reproducible even in brain regions with large susceptibility distortions. The right panel shows box plots of the ICC values in the masked regions. We can see clear drops in reproducibility from P1 to P2 acquisitions.

**Figure 8.**
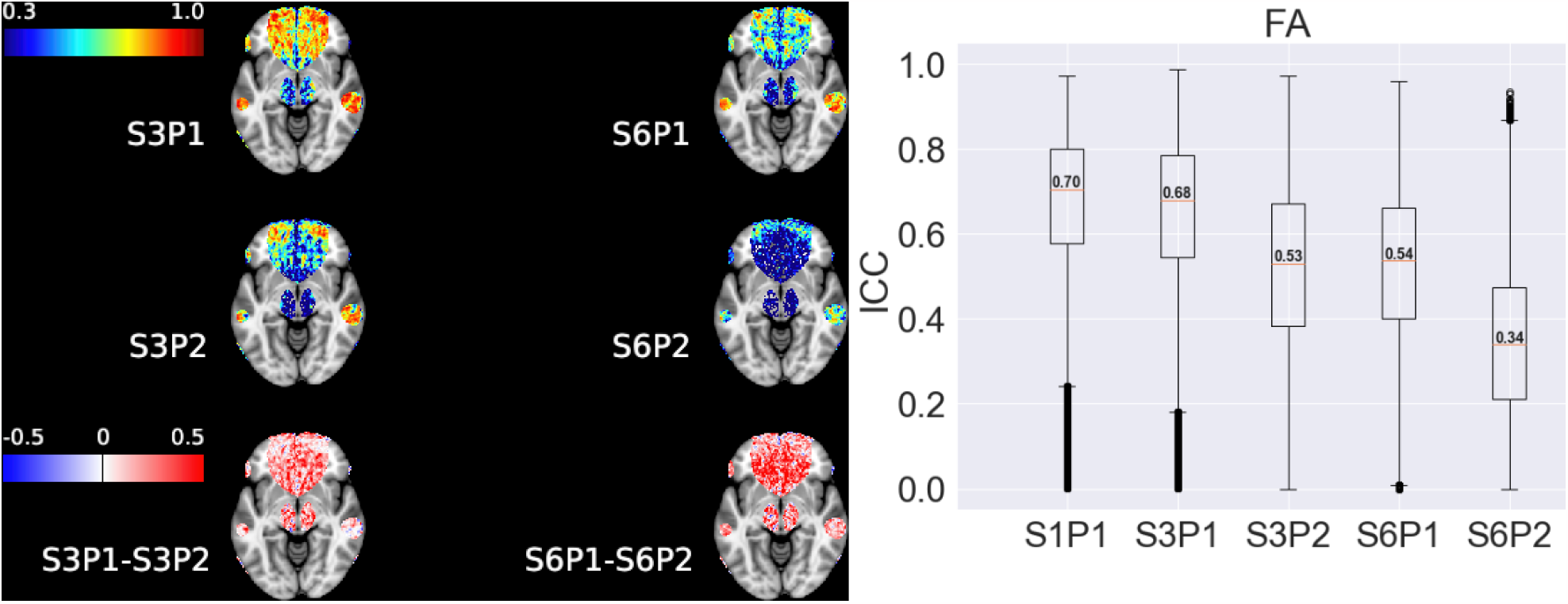
Maps of FA ICC in high-distortion areas (left) with subtraction maps shown in the bottom row. The box plot of FA ICC is shown in the right panel.

### 4.5. Impact of acceleration on group comparisons

We now examine the impact of acceleration on comparisons between MCI and healthy older adults. We focus on S1P1, S3P1, and S6P1 due to low SNR and reproducibility of S3P2 and S6P2.

#### 4.5.1. Acceleration’s impact on effect size

Figure 9 uses histogram plots to compare how the effect size magnitude (measured by the absolute value of the adjusted Cohen’s d) changes between MCI and healthy older adults across different accelerations. Recall that we focused on voxels in the WM across all S1P1, S3P1 and S6P1 acquisitions. We also marked the percentage of voxels having increased/decreased effect size magnitude compared with S1P1. From these results, we can see that:

**Figure 9.**
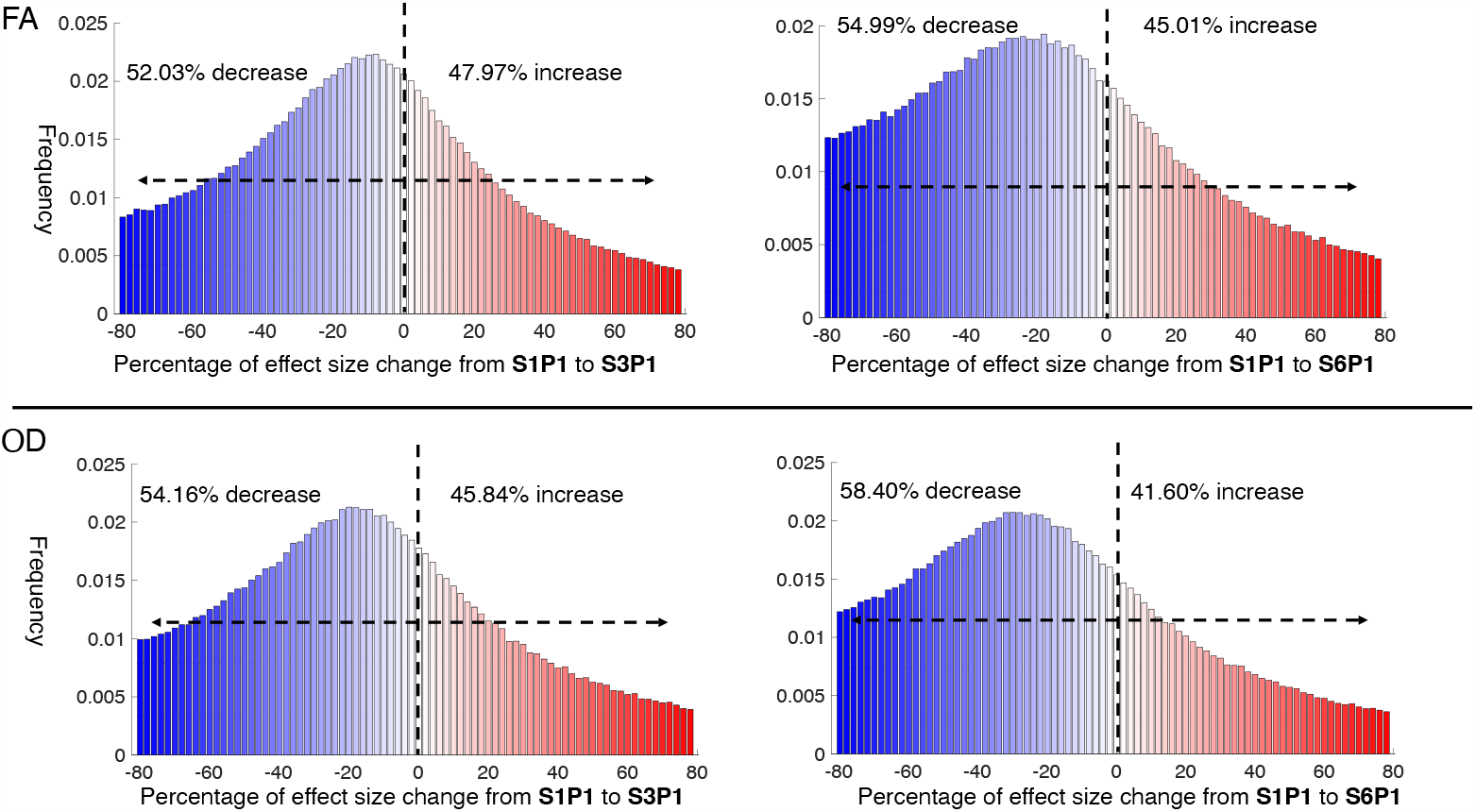
Change in effect size magnitude between accelerated and unaccelerated acquisitions for FA and OD in WM voxels. The change is measured as a percentage with respect to the magnitude of effect size at S1P1. In FA, the median effect size change was -2.52% in S3P1 and -8.11% in S6P1. In OD, the median effect size change was -5.92% in S3P1 and -13.29% in S6P1 in OD.

- In general, acceleration shrinks the effect size in more voxels than it increases it. The shrinking effect increases with acceleration. For example, for FA, from S1P1 to S3P1, 52.03% of WM voxels have decreased effect sizes while 47.97% have increased effect sizes. From S1P1 to S6P1, the proportions become 54.99% (decreased) versus 45.01% (increased).
- OD is more sensitive to acceleration than FA. 54.16% of WM voxels have a shrinking magnitude of effect size for OD at S3P1. The proportion becomes 58.40% at S6P1.

#### 4.5.2. Group comparisons in the fornix

Figure 10 shows the negative log10 p-values on the fornix template. Figure S.8 shows a similar effect size analysis as the one in Figure 9, but for voxels in the fornix. We make the following observations:

**Figure 10.**
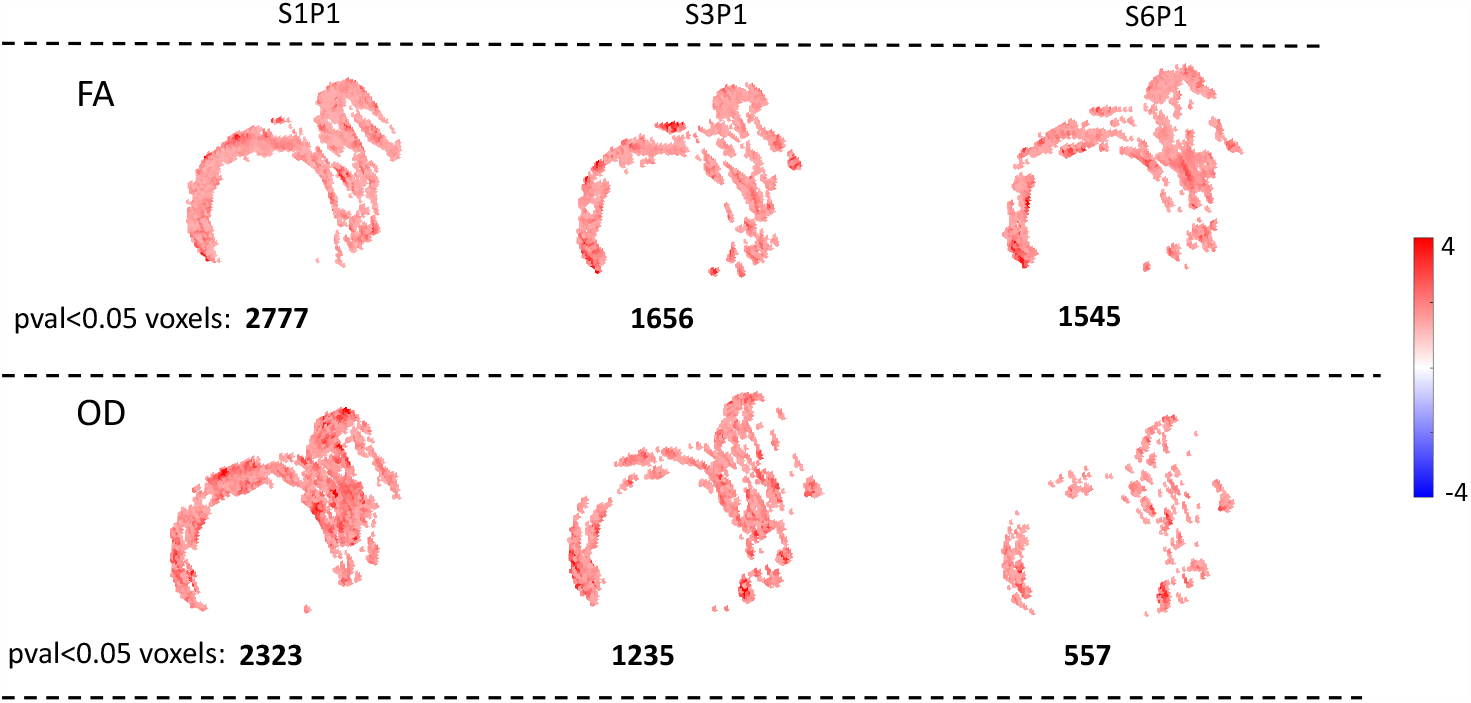
-log10(p-values) across different acceleration factors for the group difference between MCIs and healthy older adults for FA and OD in voxels located in the fornix. Results were obtained from a linear mixed model controlling for sex, age, and absolute head motion. Only voxels with p-values smaller than 0.05 are displayed (the number of voxels with p-values *<* 0.05 is displayed at the bottom of each panel).

- The fornix may be more significantly impacted by acceleration than other brain regions. The FA effect size’s amplitude was reduced from S1P1 to S3P1 in 62.8% voxels, while in the whole WM region, the number is only 52.03%. In the fornix, the median effect size of FA decreased by 13% in S3P1 and 27% in S6P1. For OD, the effect size decreased in 62.78% of voxels for S3P1 and 71.87% of voxels for S6P1, and the median effect size reduction was 13.5% for S3P1 and 28.6% for S6P1.
- For FA, S1P1 has more voxels with *p <* 0.05 than S3P1 and S6P1. There is a strong correspondence between the regions with small p-values in S1P1, S3P1, and S6P1.
- For OD, S1P1 again has more voxels with *p <* 0.05. The p-values for group differences in OD are very sensitive to acceleration, which agrees with the effect size analysis in Figure 9. The number of voxels with *p <* 0.05 in S6P1 decreased by more than 70 percent compared with S1P1.

## 5. Discussion

This study collected diffusion data from 40 older volunteers (20 MCIs) in a test-retest setting under five different acceleration factors to understand the acceleration’s impacts on WM microstructure analysis. We discuss the benefits and costs of accelerated acquisition in dMRI.

### Motion benefits

Absolute motion decreases in accelerated acquisitions. The motion was significantly reduced from S1P1 to S3P1 (Cohen’s d=-1), with additional reductions in S6P1 (Cohen’s d=-1.6) (Figure 2). The difference in motion was predictive of the differences between accelerated and unaccelerated acquisitions (Table 3 and Table S.2). Motion was more predictive in S6P1-S1P1 than in S3P1-S1P1, which coincides with greater reduction in motion for the shorter S6P1 acquisition. Previous studies on intra-subject variation in motion found that the effects of motion on FA varied by region, leading to lower FA in some regions of the corpus callosum but possible increases in the superior longitudinal fasciculus [Yendiki et al., 2014]. Our analysis examined the associations with motion across voxels on the mean data, thereby removing participant effects. Our analysis adds to the literature on motion impacts by providing evidence that the spatial variation in acceleration impacts can be partially attributed to the decrease in rotational motion in accelerated acquisitions.

### Costs of noise amplification

Noise is amplified by acceleration, which causes biases in white matter microstructure estimates. SNR and CNR progressively decrease with acceleration from S1P1 to S6P2, with particularly large costs when phase acceleration is combined with multiband acceleration (Figure 2). There is an increasing trend in FA and a decreasing trend in OD with higher acceleration factors (Figures 4 and 5). The increase in FA is consistent with previous theoretical and empirical findings that FA increases as SNR decreases at b=1000 [Bastin et al., 1998; Farrell et al., 2007; Jones and Basser, 2004]. Additionally, the CNR is more sensitive to acceleration at higher b-values (Figure 2). Since we used b=2000 in NODDI while FA used only b=1000, OD appears to be more sensitive to the biases introduced by noise amplification (Figure 5).

Since 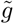-factor is spatially varying, the biases introduced by noise amplification also vary spatially. The noise amplification is spatially inhomogeneous, including large increases in WM regions near the subcallosal cortex, brainstem, and cerebellum, and less noise amplification and even signal enhancement in S=3 in some regions of the occipital lobe (Figure 3). The FA and OD differences between acquisitions vary spatially (Figures 4 and 5). This is consistent with previous findings that multiband acceleration induces spatial biases in functional connectivity studies [Risk et al., 2021].

Multiband but not phase acceleration results for reproducible microstructure metrics: We see roughly the following reproducibility order: S1P1*≈* S3P1 *>* S6P1*≥* S3P2 *»* S6P2. However, the order can differ depending on the measure and location (Figure 6). Overall, our study indicates large costs to in-plane acceleration, with large decreases in reproducibility (Figure 7). Even in regions with possible high distortion in EPI, phase acceleration decreases the reproducibility of dMRI-derived microstructure metrics (Figure 8).

### Acceleration can harm group-wise comparisons

The acceleration shrinks the effect size in more voxels than increases across all white matter voxels (Figure 9), with larger impacts in the fornix (Figure S.8). OD from the NODDI model is more sensitive to acceleration than FA derived from the DTI model. This may be due to the lower CNR at b=2000 (used in NODDI but not DTI), as higher b-values are more sensitive to noise amplification. In linear mixed models in the fornix, the p-values of MCI versus healthy control in S1P1 tended to be smaller than accelerated acquisitions, particularly for OD.

According to our findings, we have the following recommendations for studies collecting multi-shell dMRI:

1. **Moderate multiband acceleration (e.g**., **S3P1) balances costs and benefits**. Moderate acceleration can speed up the acquisition while maintaining the data’s reproducibility and the effect size between groups. Overall, S3P1 has similar reproducibility and effect size as S1P1 for the FA metric but is much faster to acquire.
2. **Aggressive multiband acceleration (e.g**., **S6P1) can be used for acquiring reproducible data**. Even S6P1 can be used to acquire white matter microstructure data, since it has good reproducibility and is much faster to acquire (about 1 min for 14 b=0 and 26 b=1000 diffusion images, about 4.5 minutes when also including 102 b=2000). It does result in reduced effect sizes.
3. **No acceleration (S1P1) should be used to achieve the biggest effect size in some brain regions**. Multiband costs on effect size are larger in the fornix than across the whole brain WM voxels. Therefore, if a study is limited by the number of participants, to gain the best statistical power, S1P1 should be used for the fornix (especially when the biomarker of interest is derived from the multishell data, e.g., OD). This can be the case for other regions that are heavily impacted by acceleration.
4. **Phase acceleration (in-plane acceleration) has large costs in dMRI**. Our results indicate that in-plane acceleration degrades dMRI-derived microstructure metrics, even in high-distortion areas. IPA in dMRI does not result in proportional decreases in acquisition time (Table 2). IPA is often used in dMRI studies to reduce spatial blurring. Our quantification of the decrease in ICC provides important information for researchers when deciding whether to use phase acceleration.

A shortcoming of this study is that we evaluated SMS factors equal to 1, 3, and 6, but other SMS factors may be optimal. SMS=3 has been popularized by the Human Connectome Project, UK Biobank, and ABCD protocols, and is thus important to include. However, previous work in fMRI suggests that SMS=4 can be optimal in cortical areas [Bhandari et al., 2020; Cahart et al., 2022]. In our previous resting-state fMRI study, we found that SMS=4 outperformed SMS=3 [Risk et al., 2021]. Unfortunately, time constraints precluded the use of both SMS=4 and SMS=6 in this study. We elected to evaluate SMS=6 because it was significantly faster than SMS=3. Since S6P1 also had good reliability in the present study, we speculate that SMS=4 with no IPA has high reliability. Additionally, there are other methods to speed up dMRI acquisition, e.g., the slice-interleaved diffusion encoding method [Xu et al., 2022], which is an important avenue for future research.

We also note that our S3P1 protocol (Table 2) is similar to SMS=3 protocols used in large-scale neuroimaging studies but with some differences. UK Biobank also used two shells and 2 mm^3^ isotropic voxels, but they collected 50 directions for each shell with a 32-channel head coil. The HCP protocol used a customized gradient coil, three shells (b=1000, b=2000, b=3000 s/mm^2^), 270 directions, 1.25 mm^3^ isotropic voxels, and a 32-channel head coil [Sotiropoulos et al., 2013]. Our protocol reflects current guidance on collecting more b=2000 than b=1000 directions due to lower SNR at b=2000, and it is a protocol that can be broadly adopted in other studies. Our use of the 64-channel head coil reflected the hardware availability at our scanner, and it is commonly used in other studies. However, it may also be a factor impacting dMRI reliability.

This study focused on white matter microstructure. An important avenue for future research is to examine the effect of acceleration on structural connectivity. In tractography studies, streamlines cross areas of high and low noise amplification, and the impacts of this spatially varying noise amplification are complex. Since we found spatially varying biases, we hypothesize acceleration will also cause biases in structural connectome studies.

## 6. Credit authorship contribution statement

**Zhengwu Zhang**: Conceptualization, Methodology, Software, Validation, Formal analysis, Investigation, Resources, Data curation, Writing-original draft, Visualization, Supervision, Project administration, Funding acquisition. **Arun Venkataraman**: Software, Validation, Formal analysis, Investigation, Resources, Data curation, Writing -original draft, Visualization. **Martin Cole**: Software, Investigation, Data curation, Visualization. **Tianrui Liu**: Investigation, Data curation, Visualization. **Deqiang Qiu**: Conceptualization, Formal analysis, Funding acquisition, Supervision. **Vankee Lin Feng**: Conceptualization, Methodology, Resources, Data curation, Writing -original draft, Supervision, Project administration, Funding acquisition. **Benjamin B. Risk**: Conceptualization, Methodology, Software, Validation, Formal analysis, Investigation, Resources, Data curation, Writing-original draft, Visualization, Supervision, Project administration, Funding acquisition.

## 7. Acknowledgments

This research was supported by NIH R21 AG066970.

## 8. Supplementary material

Supplementary material associated with this article can be found in the online version at XXXXXX.

## References

Andersson, J.L., Graham, M.S., Drobnjak, I., Zhang, H., Filippini, N., Bastiani, M., 2017. Towards a comprehensive framework for movement and distortion correction of diffusion mr images: Within volume movement. Neuroimage 152, 450–466.

Andersson, J.L., Graham, M.S., Zsoldos, E., Sotiropoulos, S.N., 2016. Incorporating outlier detection and replacement into a non-parametric framework for movement and distortion correction of diffusion MR images. NeuroImage 141, 556–572.

Andersson, J.L., Skare, S., Ashburner, J., 2003. How to correct susceptibility distortions in spin-echo echo-planar images:application to diffusion tensor imaging. NeuroImage 20, 870–888.

Andersson, J.L., Sotiropoulos, S.N., 2016. An integrated approach to correction for off-resonance effects and subject movement in diffusion MR imaging. NeuroImage 125, 1063–1078.

Avants, B.B., Tustison, N., Song, G., 2009. Advanced normalization tools (ANTS). Insight j 2, 1–35.

Bammer, R., Auer, M., Keeling, S.L., Augustin, M., Stables, L.A., Prokesch, R.W., Stollberger, R., Moseley, M.E., Fazekas, F., 2002. Diffusion tensor imaging using single-shot SENSE-EPI. Magnetic Resonance in Medicine: An Official Journal of the International Society for Magnetic Resonance in Medicine 48, 128–136.

Basser, P.J., Pierpaoli, C., 2011. Microstructural and physiological features of tissues elucidated by quantitative-diffusion-tensor mri. Journal of magnetic resonance 213, 560–570.

Bastiani, M., Cottaar, M., Fitzgibbon, S.P., Suri, S., Alfaro-Almagro, F., Sotiropoulos, S.N., Jbabdi, S., Andersson, J.L., 2019. Automated quality control for within and between studies diffusion MRI data using a non-parametric framework for movement and distortion correction. NeuroImage 184, 801–812.

Bastin, M.E., Armitage, P.A., Marshall, I., 1998. A theoretical study of the effect of experimental noise on the measurement of anisotropy in diffusion imaging. Magnetic resonance imaging 16, 773–785.

Baum, G.L., Roalf, D.R., Cook, P.A., Ciric, R., Rosen, A.F., Xia, C., Elliott, M.A., Ruparel, K., Verma, R., Tunç, B., et al., 2018. The impact of in-scanner head motion on structural connectivity derived from diffusion MRI. Neuroimage 173, 275–286.

Bhandari, R., Kirilina, E., Caan, M., Suttrup, J., De Sanctis, T., De Angelis, L., Keysers, C., Gazzola, V., 2020. Does higher sampling rate (multiband+SENSE) improve group statistics - An example from social neuroscience block design at 3T. NeuroImage 213, 116731.

Bouyagoub, S., Dowell, N.G., Gabel, M., Cercignani, M., 2021. Comparing multiband and singleband EPI in NODDI at 3 T: what are the implications for reproducibility and study sample sizes? Magnetic Resonance Materials in Physics, Biology and Medicine 34, 499–511.

Breuer, F.A., Kannengiesser, S.A., Blaimer, M., Seiberlich, N., Jakob, P.M., Griswold, M.A., 2009. General formulation for quantitative G-factor calculation in GRAPPA reconstructions. Magnetic Resonance in Medicine 62, 739–746.

Brown, C.A., Johnson, N.F., Anderson-Mooney, A.J., Jicha, G.A., Shaw, L.M., Trojanowski, J.Q., Van Eldik, L.J., Schmitt, F.A., Smith, C.D., Gold, B.T., 2017. Development, validation and application of a new fornix template for studies of aging and preclinical Alzheimer’s disease. NeuroImage: Clinical 13, 106–115.

Cahart, M.S., O’Daly, O., Giampietro, V., Timmers, M., Streffer, J., Einstein, S., Zelaya, F., Dell’Acqua, F., Williams, S.C.R., 2022. Comparing the test–retest reliability of resting-state functional magnetic resonance imaging metrics across single band and multiband acquisitions in the context of healthy aging. Human Brain Mapping 44, 1901–1912.

Casey, B.J., Cannonier, T., Conley, M.I., Cohen, A.O., Barch, D.M., Heitzeg, M.M., Soules, M.E., Teslovich, T., Dellarco, D.V., Garavan, H., et al., 2018. The adolescent brain cognitive development (ABCD) study: imaging acquisition across 21 sites. Developmental Cognitive Neuroscience 32, 43–54.

Demetriou, L., Kowalczyk, O.S., Tyson, G., Bello, T., Newbould, R.D., Wall, M.B., 2018. A comprehensive evaluation of increasing temporal resolution with multiband-accelerated protocols and effects on statistical outcome measures in fMRI. NeuroImage 176, 404–416.

Descoteaux, M., Wiest-Daesslé, N., Prima, S., Barillot, C., Deriche, R., 2008. Impact of Rician adapted non-local means filtering on HARDI, in: International Conference on Medical Image Computing and Computer-Assisted Intervention, Springer. pp. 122–130.

Duan, F., Zhao, T., He, Y., Shu, N., 2015. Test–retest reliability of diffusion measures in cerebral white matter: A multiband diffusion MRI study. Journal of Magnetic Resonance Imaging 42, 1106–1116.

Farrell, J.A., Landman, B.A., Jones, C.K., Smith, S.A., Prince, J.L., Van Zijl, P.C., Mori, S., 2007. Effects of signal-to-noise ratio on the accuracy and reproducibility of diffusion tensor imaging–derived fractional anisotropy, mean diffusivity, and principal eigenvector measurements at 1.5 t. Journal of Magnetic Resonance Imaging: An Official Journal of the International Society for Magnetic Resonance in Medicine 26, 756–767.

Feinberg, D.A., Setsompop, K., 2013. Ultra-fast MRI of the human brain with simultaneous multi-slice imaging. Journal of Magnetic Resonance 229, 90–100.

Garyfallidis, E., Brett, M., Amirbekian, B., Rokem, A., Van Der Walt, S., Descoteaux, M., Nimmo-Smith, I., 2014. Dipy, a library for the analysis of diffusion MRI data. Frontiers in Neuroinformatics 8, 8.

Griswold, M.A., Jakob, P.M., Heidemann, R.M., Nittka, M., Jellus, V., Wang, J., Kiefer, B., Haase, A., 2002. Generalized autocalibrating partially parallel acquisitions (GRAPPA). Magnetic Resonance in Medicine: An Official Journal of the International Society for Magnetic Resonance in Medicine 47, 1202–1210.

Harms, M., Somerville, L., Ances, B., Andersson, J., Barch, D., Bastiani, M., Yacoub, E., 2018. Imaging in the human connectome projects in development and aging: connectomics across the lifespan. Neuroimage 183, 972–984.

Jones, D.K., Basser, P.J., 2004. “squashing peanuts and smashing pumpkins”: how noise distorts diffusion-weighted mr data. Magnetic Resonance in Medicine: An Official Journal of the International Society for Magnetic Resonance in Medicine 52, 979–993.

Le Bihan, D., 2003. Looking into the functional architecture of the brain with diffusion MRI. Nature Reviews Neuroscience 4, 469–480.

Le Bihan, D., Breton, E., Lallemand, D., Grenier, P., Cabanis, E., Laval-Jeantet, M., 1986. MR imaging of intravoxel incoherent motions: application to diffusion and perfusion in neurologic disorders. Radiology 161, 401–407.

Lee, K., Wild, J., Griffiths, P., Paley, M., 2005. Simultaneous multislice imaging with slice-multiplexed RF pulses. Magnetic Resonance in Medicine: An Official Journal of the International Society for Magnetic Resonance in Medicine 54, 755–760.

Lucignani, M., Breschi, L., Espagnet, M.C.R., Longo, D., Talamanca, L.F., Placidi, E., Napolitano, A., 2021. Reliability on multiband diffusion NODDI models: a test retest study on children and adults. NeuroImage 238, 118234.

Merboldt, K.D., Hanicke, W., Frahm, J., 1985. Self-diffusion NMR imaging using stimulated echoes. Journal of Magnetic Resonance (1969) 64, 479–486.

Miller, K.L., Alfaro-Almagro, F., Bangerter, N.K., Thomas, D.L., Yacoub, E., Xu, J., Bartsch, A.J., Jbabdi, S., Sotiropoulos, S.N., Andersson, J.L., et al., 2016. Multimodal population brain imaging in the UK Biobank prospective epidemiological study. Nature Neuroscience 19, 1523–1536.

Moeller, S., Yacoub, E., Olman, C.A., Auerbach, E., Strupp, J., Harel, N., Ugurbil, K., 2010. Multiband multislice GE-EPI at 7 tesla, with 16-fold acceleration using partial parallel imaging with application to high spatial and temporal whole-brain fMRI. Magnetic Resonance in Medicine 63, 1144–1153.

Moseley, M.E., Kucharczyk, J., Asgari, H.S., Norman, D., 1991. Anisotropy in diffusion-weighted mri. Magnetic Resonance in Medicine 19, 321–326.

Nooner, K.B., Colcombe, S.J., Tobe, R.H., Mennes, M., Benedict, M.M., Moreno, A.L., Panek, L.J., Brown, S., Zavitz, S.T., Li, Q., et al., 2012. The NKI-Rockland sample: a model for accelerating the pace of discovery science in psychiatry. Frontiers in Neuroscience 6, 152.

Oishi, K., Zilles, K., Amunts, K., Faria, A., Jiang, H., Li, X., Akhter, K., Hua, K., Woods, R., Toga, A.W., et al., 2008. Human brain white matter atlas: identification and assignment of common anatomical structures in superficial white matter. Neuroimage 43, 447–457.

Park, H.J., Friston, K., 2013. Structural and functional brain networks: from connections to cognition. Science 342, 1238411.

Pasternak, O., Kelly, S., Sydnor, V.J., Shenton, M.E., 2018. Advances in microstructural diffusion neuroimaging for psychiatric disorders. Neuroimage 182, 259–282.

Preibisch, C., Bührer, M., Riedl, V., others, 2015. Evaluation of multiband EPI acquisitions for resting state fMRI. PloS one 10, e0136961.

Pruessmann, K.P., Weiger, M., Scheidegger, M.B., Boesiger, P., 1999. SENSE: sensitivity encoding for fast MRI. Magnetic Resonance in Medicine: An Official Journal of the International Society for Magnetic Resonance in Medicine 42, 952–962.

Risk, B., Kociuba, M., Rowe, D., 2018. Impacts of simultaneous multislice acquisition on sensitivity and specificity in fMRI. NeuroImage 172.

Risk, B.B., Murden, R.J., Wu, J., Nebel, M.B., Venkataraman, A., Zhang, Z., Qiu, D., 2021. Which multiband factor should you choose for your resting-state fMRI study? NeuroImage 234, 117965.

Shrout, P.E., Fleiss, J.L., 1979. Intraclass correlations: uses in assessing rater reliability. Psychological Bulletin 86, 420.

Sotiropoulos, S.N., Jbabdi, S., Xu, J., Andersson, J.L., Moeller, S., Auerbach, E.J., Glasser, M.F., Hernandez, M., Sapiro, G., Jenkinson, M., et al., 2013. Advances in diffusion MRI acquisition and processing in the Human Connectome Project. Neuroimage 80, 125–143.

Srirangarajan, T., Mortazavi, L., Bortolini, T., Moll, J., Knutson, B., 2021. Multi-band FMRI compromises detection of mesolimbic reward responses. NeuroImage 244, 118617.

Tax, C.M., Grussu, F., Kaden, E., Ning, L., Rudrapatna, U., Evans, C.J., St-Jean, S., Leemans, A., Koppers, S., Merhof, D., 2019. Cross-scanner and cross-protocol diffusion MRI data harmonisation: A benchmark database and evaluation of algorithms. Neuroimage 195, 285–299.

Taylor, D., Bushell, M., 1985. The spatial mapping of translational diffusion coefficients by the NMR imaging technique. Physics in Medicine & Biology 30, 345.

Todd, N., Josephs, O., Zeidman, P., Flandin, G., Moeller, S., Weiskopf, N., 2017. Functional sensitivity of 2D simultaneous multi-slice echo-planar imaging: effects of acceleration on g-factor and physiological noise. Frontiers in Neuroscience 11.

Tustison, N.J., Avants, B.B., Cook, P.A., Zheng, Y., Egan, A., Yushkevich, P.A., Gee, J.C., 2010. N4ITK: improved N3 bias correction. IEEE Transactions on Medical Imaging 29, 1310–1320.

Uğurbil, K., Xu, J., Auerbach, E.J., Moeller, S., Vu, A.T., Duarte-Carvajalino, J.M., Lenglet, C., Wu, X., Schmitter, S., Van de Moortele, P.F., et al., 2013. Pushing spatial and temporal resolution for functional and diffusion MRI in the Human Connectome Project. NeuroImage 80, 80–104.

Weiner, M.W., Veitch, D.P., Aisen, P.S., Beckett, L.A., Cairns, N.J., Green, R.C., Harvey, D., Jack Jr, C.R., Jagust, W., Morris, J.C., et al., 2017. The Alzheimer’s Disease Neuroimaging Initiative 3: Continued innovation for clinical trial improvement. Alzheimer’s & Dementia 13, 561–571.

Xu, T., Wu, Y., Hong, Y., Ahmad, S., Huynh, K.M., Wang, Z., Lin, W., Chang, W.T., Yap, P.T., 2022. Rapid diffusion magnetic resonance imaging using slice-interleaved encoding. Medical Image Analysis 81, 102548.

Yendiki, A., Koldewyn, K., Kakunoori, S., Kanwisher, N., Fischl, B., 2014. Spurious group differences due to head motion in a diffusion mri study. Neuroimage 88, 79–90.

Zhang, H., Schneider, T., Wheeler-Kingshott, C.A., Alexander, D.C., 2012. NODDI: practical in vivo neurite orientation dispersion and density imaging of the human brain. NeuroImage 61, 1000–1016.

Zhao, T., Duan, F., Liao, X., Dai, Z., Cao, M., He, Y., Shu, N., 2015. Test-retest reliability of white matter structural brain networks: a multiband diffusion MRI study. Frontiers in Human Neuroscience 9, 59.

