## Supplementary material for "The impact of multiband and in-plane acceleration on white matter microstructure analysis"

### 1. Supplementary Figures

Table S.1: Effect sizes of QC metrics between P1 and P2 acquisitions.

| Effect sizes of QC metrics | | | | |
|---|---|---|---|---|
| Groups | Absolute Motion | SNR b=0 | CNR b=1000 | CNR b=2000 |
| S3P2 - S3P1 | −0.32 | −0.67 | −2.86 | −3.71 |
| S6P2 - S6P1 | −0.46 | −3.42 | −7.21 | −8.03 |

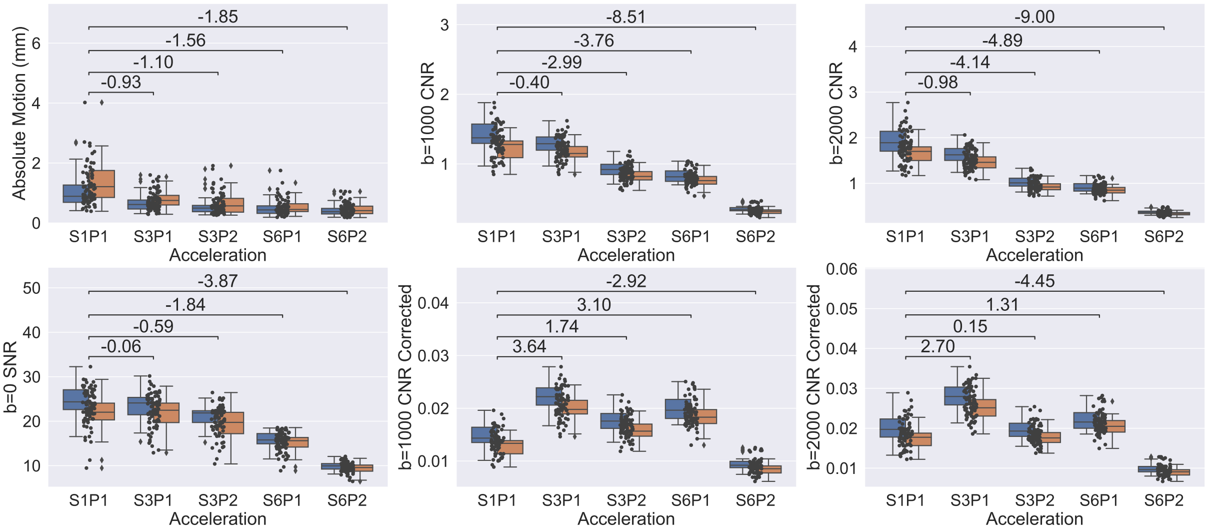


Figure S.1: QC metrics stratified by healthy and MCI participants of absolute motion (left in row one), b=0 SNR (left in row two), CNR and corrected CNR of $b = 1000$ shell (middle column), and CNR and corrected CNR of $b = 2000$ shell (right column). The corrected CNR is normalized CNR with respect to the repetition time (i.e., CNR/$\sqrt{TR}$). The numbers above bars represent Cohen's D effect size calculated for the MCI subjects.

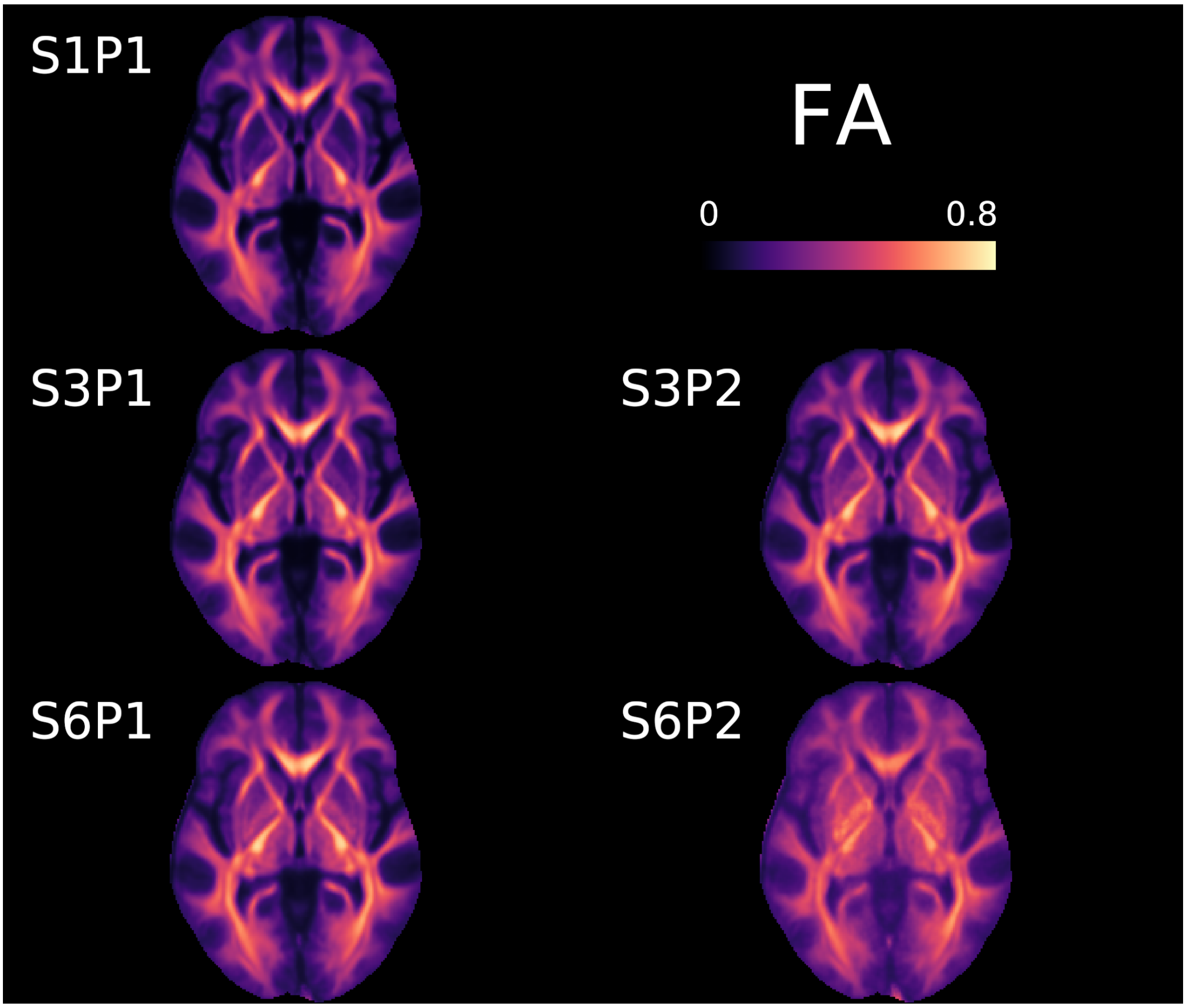


Figure S.2: FA averaged across all scans. Similar to Figure 6 in the main manuscript but uses a different color bar (0 to 0.8).

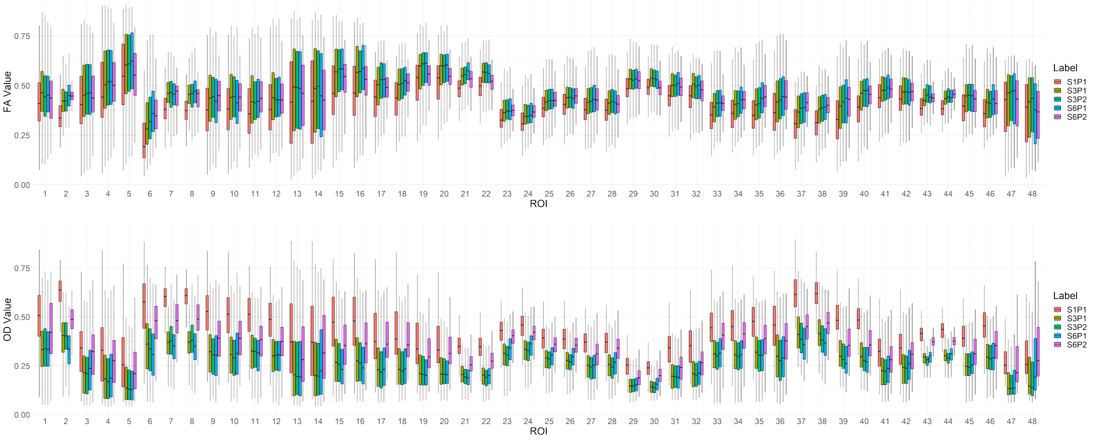


Figure S.3: Box plots of FA (top) and OD (bottom) averaged across all scans in all 48 ROIs in the ICBM-DTI-81 white-matter labels atlas.

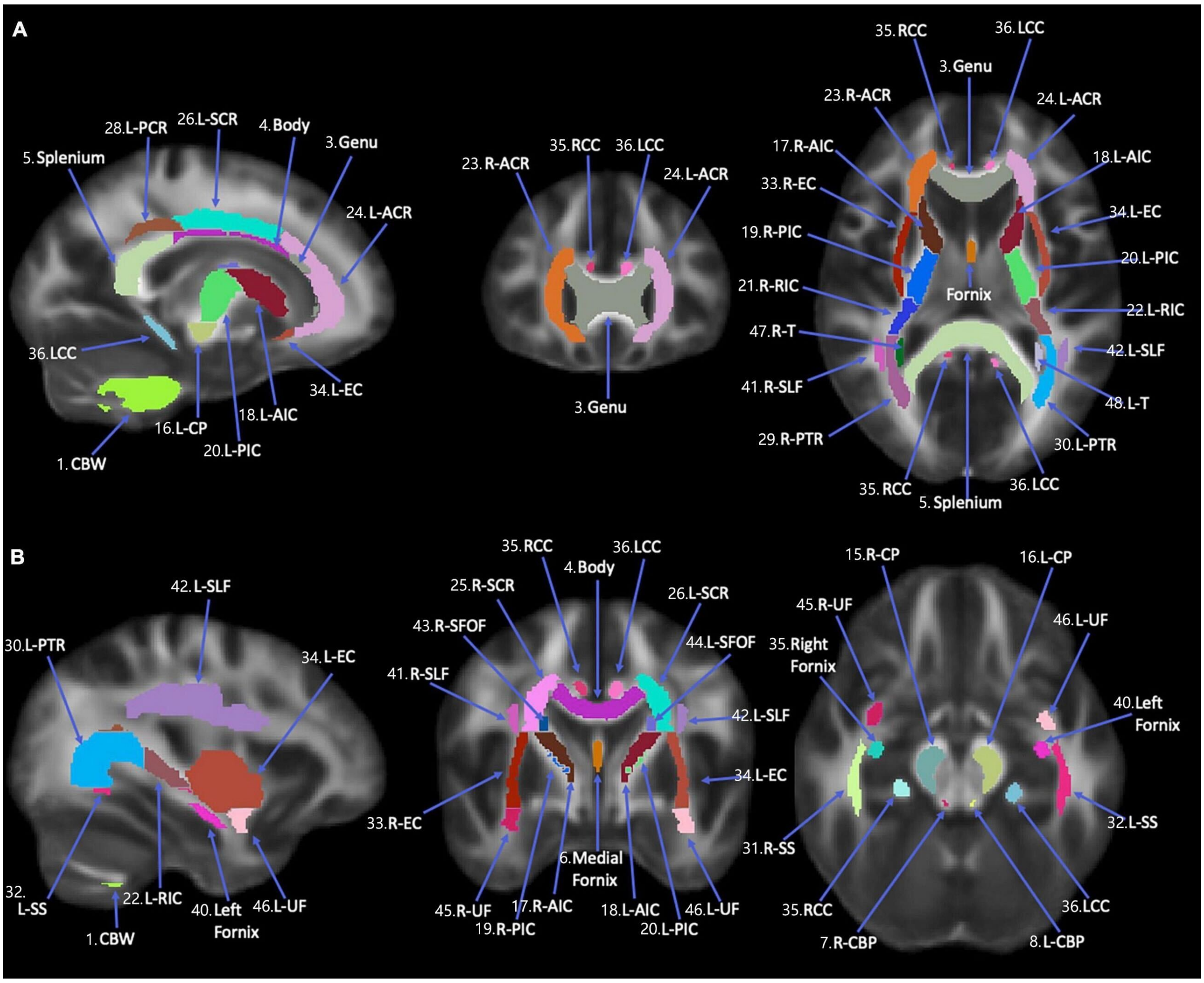


Figure S.4: The location of 48 ROIs in the ICBM-DTI-81 white-matter labels atlas.

Table S.2: Performance for predicting FA and OD bias using $\tilde{g}$-factor ($x_1$) and motion difference ($x_2$) for phase accelerated images. Each entry shows the correlation between predicted and measured values in an independent test dataset.

| | S3P2-S1P1 FA | | | | S6P2-S1P1 FA | | | |
|---|---|---|---|---|---|---|---|---|
| Model | $(x_1, x_2)$ | $(x_1, x_2, \log(x_2))$ | $x_1$ | $x_2$ | $(x_1, x_2)$ | $(x_1, x_2, \log(x_2))$ | $x_1$ | $x_2$ |
| Linear Regression | 0.50 | 0.50 | 0.50 | 0.32 | 0.18 | 0.23 | 0.14 | −0.03 |
| GAM | 0.49 | 0.51 | 0.49 | 0.34 | 0.17 | 0.25 | 0.13 | −0.01 |
| Random Forest | 0.45 | 0.45 | 0.37 | 0.24 | 0.15 | 0.15 | 0.03 | 0.00 |
| XGBoost | 0.46 | 0.46 | 0.49 | 0.34 | 0.20 | 0.20 | 0.13 | −0.03 |
| SVM | 0.50 | 0.51 | 0.50 | 0.34 | 0.21 | 0.18 | 0.11 | −0.03 |
| Gradient Boosting | 0.50 | 0.50 | 0.48 | 0.34 | 0.18 | 0.18 | 0.11 | −0.02 |
| | S3P2-S1P2 OD | | | | S6P2-S1P1 OD | | | |
| Linear Regression | 0.55 | 0.55 | 0.55 | 0.35 | 0.05 | 0.12 | 0.01 | 0.05 |
| GAM | 0.54 | 0.52 | 0.56 | 0.35 | 0.22 | 0.23 | 0.24 | 0.08 |
| Random Forest | 0.45 | 0.45 | 0.44 | 0.21 | 0.18 | 0.18 | 0.12 | 0.03 |
| XGBoost | 0.47 | 0.47 | 0.56 | 0.34 | 0.22 | 0.22 | 0.24 | 0.07 |
| SVM | 0.48 | 0.49 | 0.56 | 0.23 | 0.22 | 0.21 | 0.24 | 0.06 |
| Gradient Boosting | 0.55 | 0.55 | 0.55 | 0.33 | 0.22 | 0.22 | 0.23 | 0.06 |

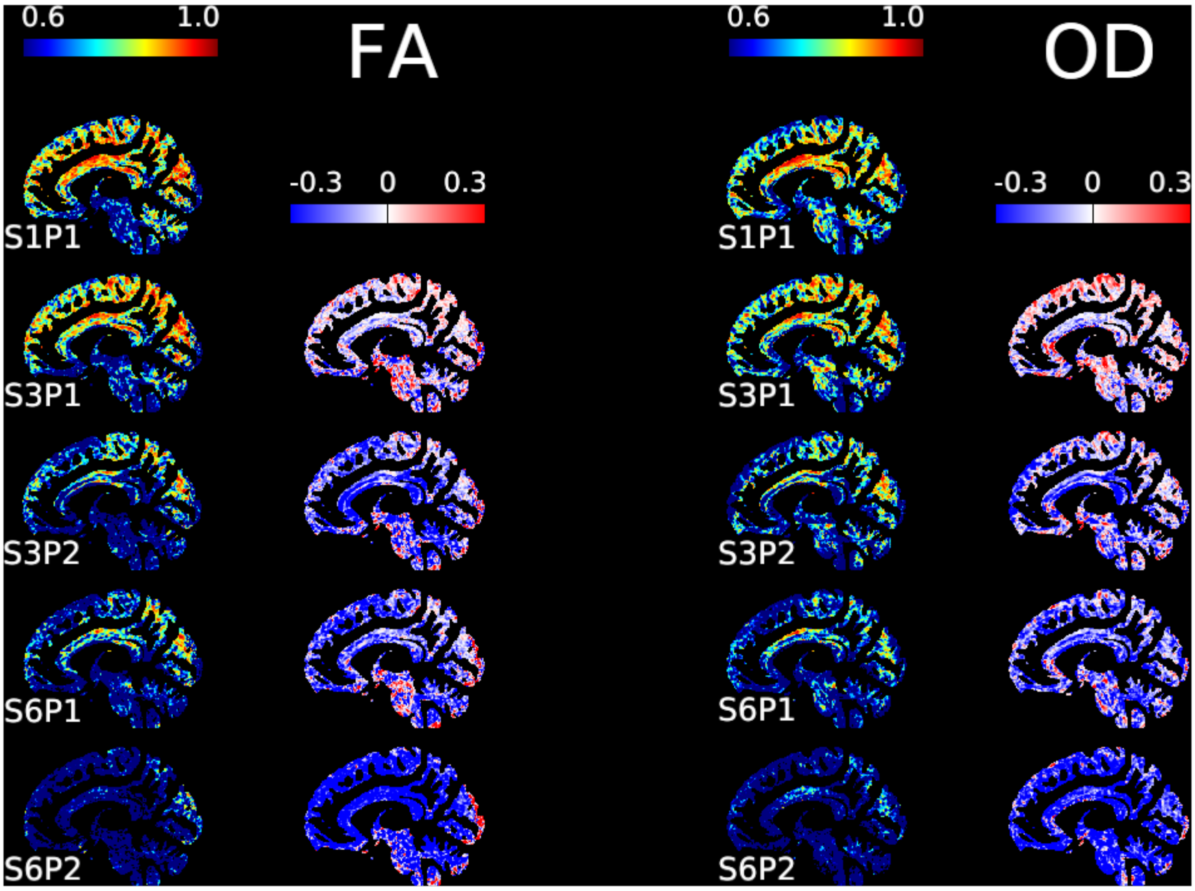


Figure S.5: ICC values of FA (left) and OD (right) calculated from 20 MCI participants.

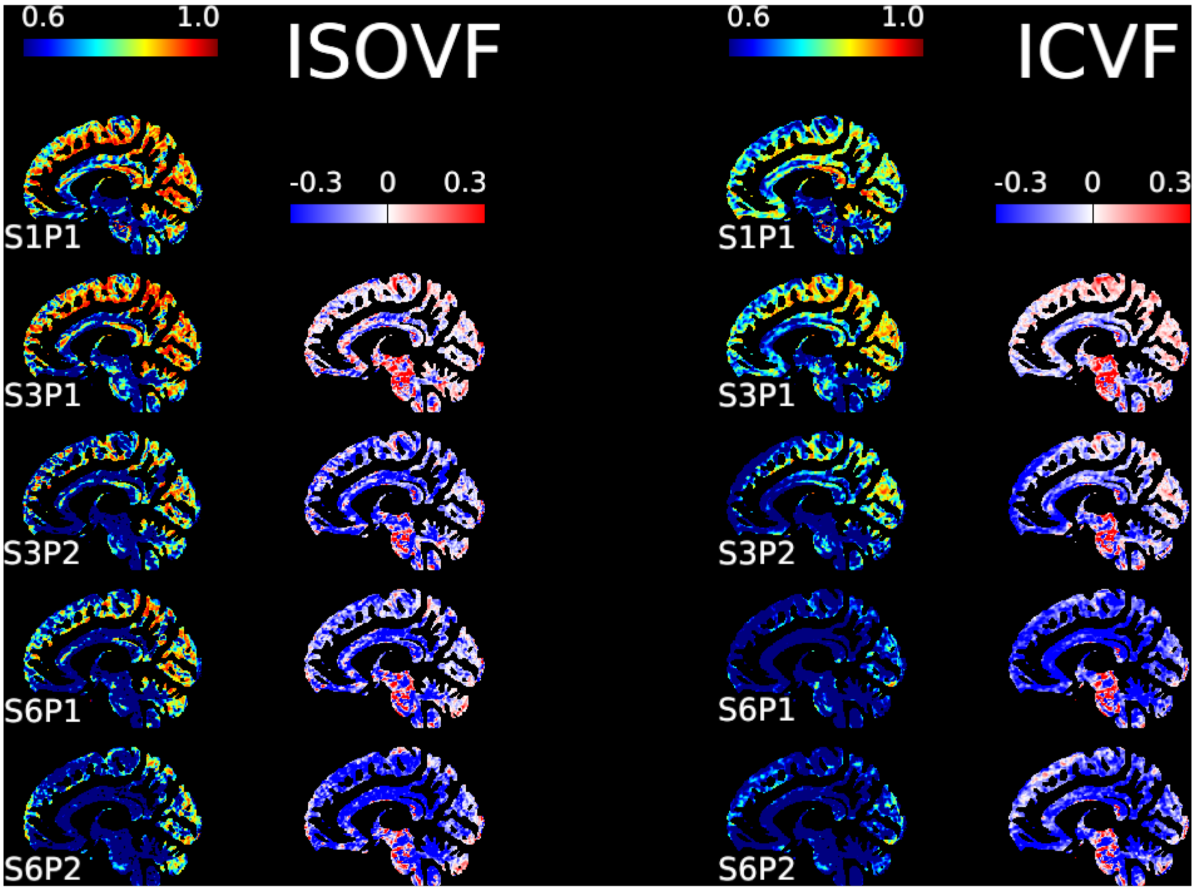


Figure S.6: Similar to Figure 8, for NODDI metrics ICVF and ISOVF.

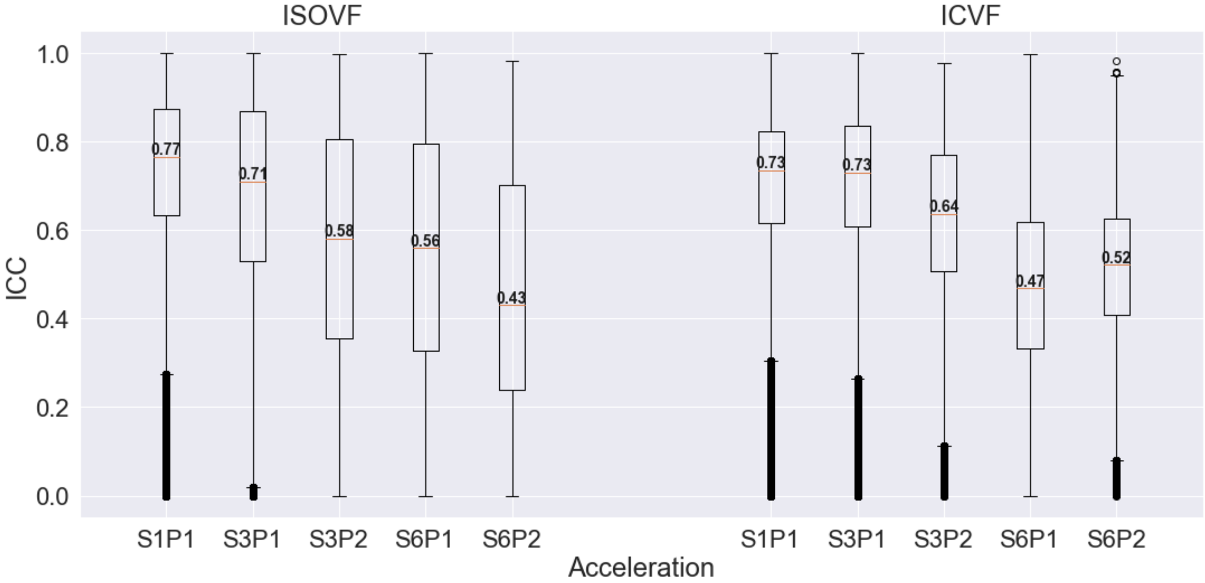


Figure S.7: Similar to Figure 9, for NODDI metrics ICVF and ISOVF.

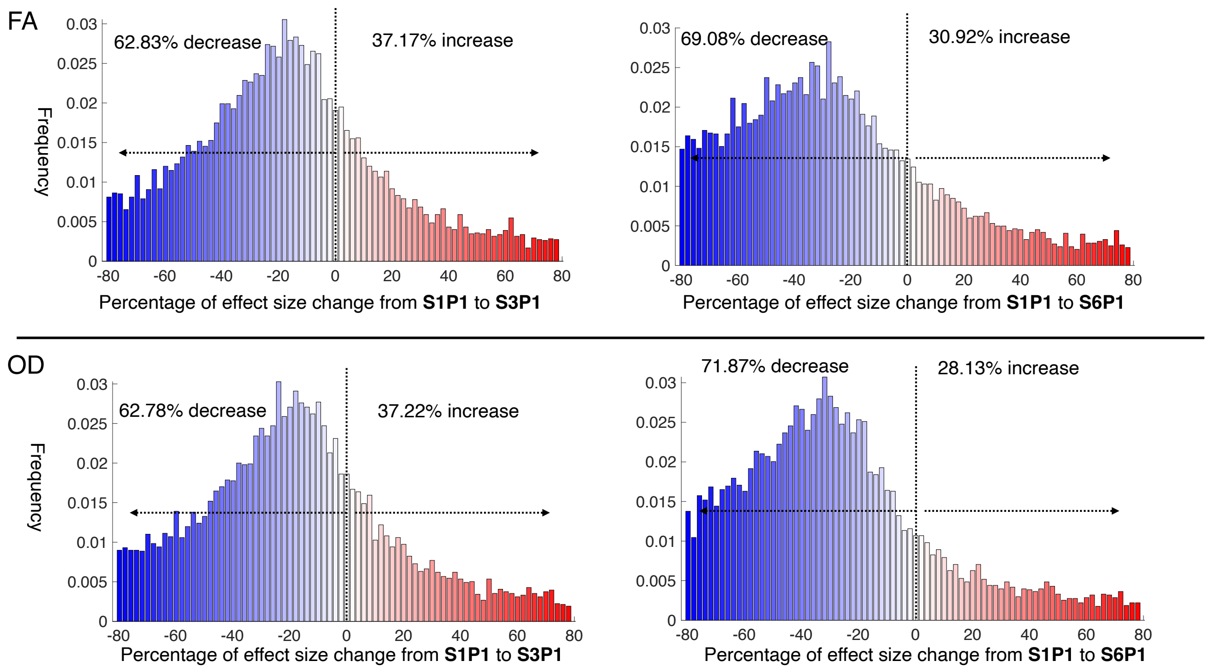


Figure S.8: The change of effect size magnitude across acquisitions for FA and OD in the fornix. In each voxel, the change is measured as a percentage of the magnitude of effect size at S1P1.
